# Integrated Ligand and Structure based approaches towards developing novel Janus Kinase 2 inhibitors for the treatment of myeloproliferative neoplasms

**DOI:** 10.1101/2020.11.26.399907

**Authors:** Unni.P Ambili, Girinath G. Pillai, Lulu.S Sajitha

## Abstract

Myeloproliferative neoplasms (MPNs) are a group of diseases affecting hematopoiesis in humans. Types of MPNs include Polycythemia Vera (PV), Essential Thrombocythemia (ET) and myelofibrosis. JAK2 gene mutation at 617^th^ position act as a major causative factor for the onset and progression of MPNs. So, JAK2 inhibitors are widely used for the treatment of MPNs. But, increased incidence of adverse drug reactions associated with JAK2 inhibitors acts as a paramount challenge in the treatment of MPNs. Hence, there exists an urgent need for the identification of novel lead molecules with enhanced potency and bioavailability. We employed ligand and structure-based approaches to identify novel lead molecules which could act as JAK2 inhibitors. The dataset for QSAR modeling (ligand-based approach) comprised of 49 compounds. We have developed a QSAR model, which has got statistical as well as biological significance. Further, all the compounds in the dataset were subjected to molecular docking and bioavailability assessment studies. Derivative compounds with higher potency and bioavailability were identified for the best lead molecule present in the dataset by employing chemical space exploration. Dataset and models are available at https://github.com/giribio/agingdata

Graphical abstract

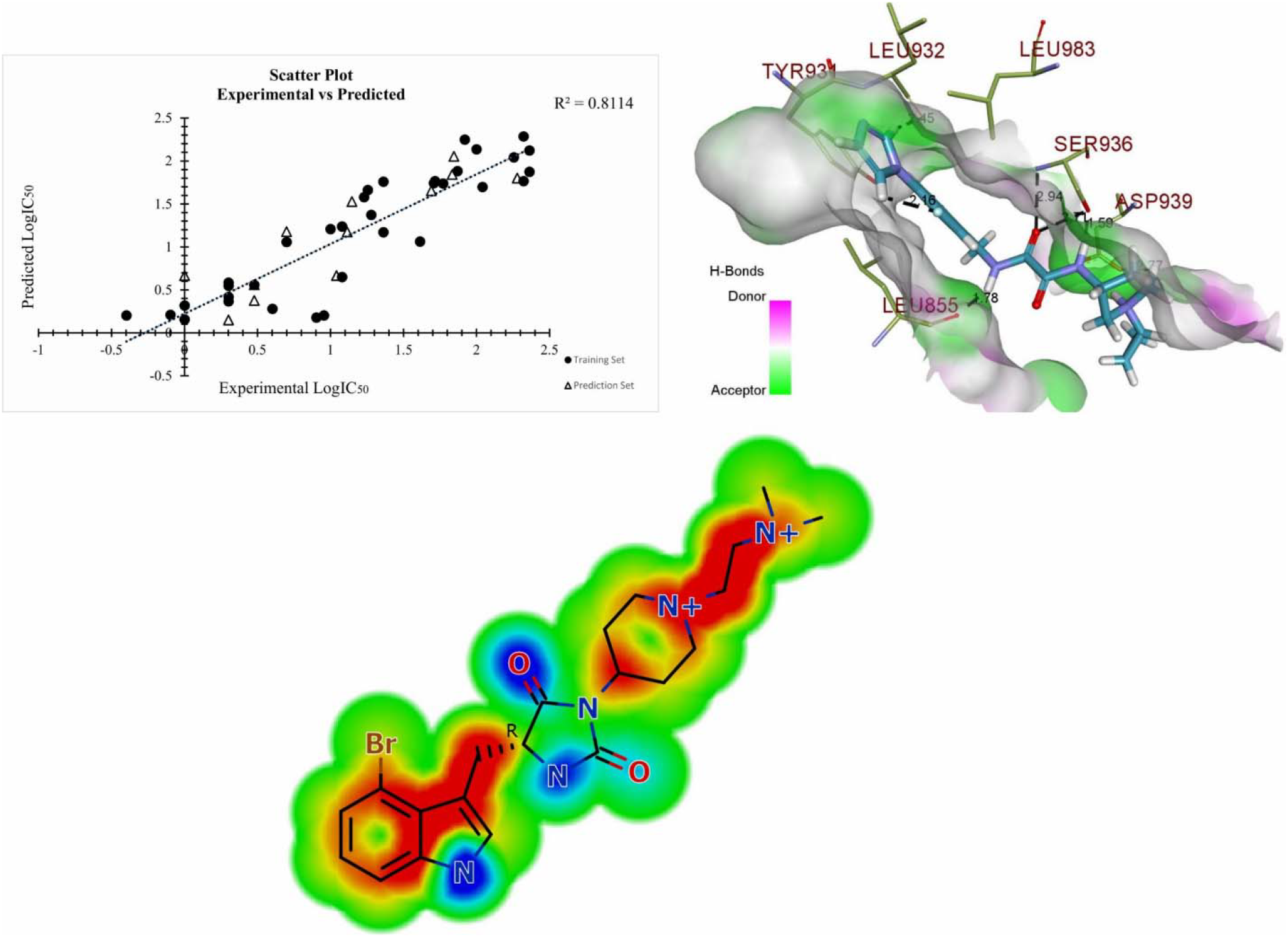

## 1. Introduction

Myeloproliferative neoplasms are a group of diseases associated with the enhanced production of blood cells from bone marrow (Hermouet et al., 2015). MPNs are classified into three groups namely Polycythemia Vera (PV), Essential Thrombocythemia (ET) and Primary Myelofibrosis (PMF) (Swerdlow et al., 2017). PV is associated with the increased count of erythrocytes and an abnormal increase in the number of cells in megakaryocytic/granulocytic lineages. ET accounts for the increase in the number of platelets while PMF is characterized by the presence of bone marrow fibrosis and enhanced cell count of megakaryocytes (Aaron T. Gerds, 2016). Mutations prevalent in in Hematopoietic Stem Cells (HSC) are considered as the major causative factor for the onset of MPNs (Mead and Mullally, 2017). Mutations responsible for the onset and progression of MPNs include Janus Kinase 2 (JAK2) gene mutations, thrombopoietin receptor gene (MPL) mutations, mutations in the calreticulin gene (CALR) and somatic mutations in the exon 2 of LNK (Shammo and Stein, 2016).

JAK2V617F is the first recognized mutation inherent to the MPN population. JAK2V617F mutation is one of the most recurrent mutations found in the clonal hematopoiesis related to aging (Steensma et al., 2015). JAK2V617F mutation is present in 95-97% of patients suffering from PV while 50-60% of patients possess JAK2V617F mutation in ET and PMF conditions (Baxter et al., 2005).

JAK2 is a non-receptor tyrosine kinase that belongs to the Janus kinase family. The activation of JAK2 protein mediated by tyrosine phosphorylation is responsible for the regulation of cytokine signaling pathways. JAK2 gene possesses an active tyrosine kinase domain called as JAK homology 1(JH1) domain, pseudokinase domain, JAK homology 2 domain (JH2), SRC homology domain (SH2) and an amino-terminal FERM domain (Ghoreschi et al., 2009). The upper surface of the N-terminal lobe of the JH2 domain is characterized by the presence of highly conserved valine at 617^th^ position. JAK2V617F mutation is associated with the substitution of phenylalanine at 617^th^ position. This mutation accounts for the destabilization of three-dimensional structure of the JH2 domain and leads to aberrant activation of JAK2 (Levine et al., 2005). The constitutive activation of JAK2 leads to the elevated production of proinflammatory cytokines, which results in immune dysregulation (Mondet et al., 2015).

The enhanced production of proinflammatory cytokines is a characteristic feature of MPNs. Neurohormonal stimulatory factors as well, as cellular responses play a significant role in regulating inflammatory cascade in healthy individuals. But, individuals affected with MPNs exhibit dysregulation of this system and hence contribute to the elevated production of proinflammatory cytokines. Higher levels of circulating inflammatory cytokines contribute to the onset of chronic inflammation and several other disease conditions. The co-morbidities associated with chronic inflammation in MPN patients include atherosclerosis, the onset of hematological and non-hematological secondary cancer (Frederiksen et al., 2011) and thrombotic events (Barbui et al., 2013) and Rheumatoid Arthritis (RA) (Barbui et al., 2013). Hence, an increased prevalence of JAK2-V617F mutations in the elder population not only contributes to the development of MPNs but also cardiovascular and other chronic inflammatory diseases.

The discovery of specific JAK2 mutation as the major contributing factor for the onset of MPNs directed towards the development of JAK2 inhibitors, which could be used as potential therapeutic agents (Reddy et al., 2012). Compounds belonging to different chemical classes such as pyrazines, pyrimidines, azaindoles, aminoindazoles, deazapurines, stilbenes, benzoxazoles , and quinoxalines were studied for their potency and specificity towards JAK2 inhibition (Baskin et al., 2010).

Hence, our study on the development of JAK2 inhibitors possessing improved pharmacokinetic features could be a promising approach for treating MPNs and co-morbidities.

## 2. Materials and methods

The objective of the present work is to identify the physicochemical properties of compounds possessing JAK2 inhibitory activity by employing SAR modeling and to predict new chemical compounds with better efficacy and pharmacokinetic properties. This objective is accomplished in two stages. 1. By building the QSAR model for compounds reported to possess JAK2 inhibitory property and predicting the activity of new chemicals using validated QSAR model. 2. Identification of binding efficacy and pharmacokinetic properties for compounds used in QSAR model building and for predicted derivatives.

### 2.1 Dataset construction

In this current research work, 49 compounds possessing JAK2 inhibitory activity were subjected to QSAR study (J. et al., 2011). The JAK2 inhibitory values reported in a negative logarithmic scale (pIC_50_) were used as the response variable for QSAR modeling. The molecular structures of chemical compounds were drawn in Marvin Sketch. 2D coordinates of chemical compounds sketched in Marvin Sketch were converted into 3D co-ordinates after the addition of explicit hydrogens. Geometry optimization of 3D co-ordinates was performed by MOPAC using PM7 parameterization (Stewart, 1989).

### 2.2 Descriptor calculation and dataset classification

Molecular descriptors were calculated by CodessaPro and PaDELsoftwares. CodessaPro is a software platform that calculates constitutional, topological, geometrical, electrostatic, quantum chemical and thermodynamic descriptors (Katritzky et al., 2001) while PaDEL is a standalone software platform, that calculates molecular descriptors and fingerprints using the Chemistry Development Kit (Yap, 2011). MOPAC optimized geometries were employed for generating descriptors from CodessaPro and PaDEL by retaining the optimized 3D configuration of chemical compounds. Descriptors generated from both softwares were combined manually without redundancy for attaining wide coverage of descriptors for QSAR modeling. Dataset of 49 compounds were classified into training and test set based on structure-based clustering analysis. Clustering was executed by StarDrop by fixing Tanimoto similarity coefficient at 0.8. The selection of test set compounds was made by considering the diversity in structure and wide range of activity within the dataset.

### 2.3 Model building

QSAR model for the set of 49 compounds was generated in QSARINS, which employs the Ordinary Least Squares method and Genetic Algorithm (GA) for feature selection (Gramatica et al., 2013). The biological activity is correlated with the physicochemical properties of compounds by Multiple Linear Regression equation,

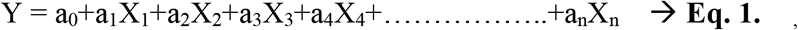

Where Y represents biological activity, X represents molecular descriptor types and a_n_ represents descriptor coefficients.

The negative logarithmic values of biological activity were set as dependent/response variable while calculated descriptors were designated as the independent variable. Feature selection was carried out by GA for 10,000 iterations. The mutation rate and population size were fixed at 20 and 10 respectively.

### 2.4 Model validation

Internal and external validation was performed for the generated QSAR model. These validation strategies ensure the reliability and accuracy of the model prediction.

#### 2.4.1 Internal validation

The predictive capacity of the QSAR model is established by considering internal and external validation procedures (Cherkasov et al., 2014). Internal validation was carried out by considering the data which created the model, while external validation was carried out by using the data which were not used for QSAR model building (Majumdar and Basak, 2018). The fitness of the model is reflected in the value of r^2^. But higher value of r^2^ does not guarantee of physical relevance, predictive capacity, and robustness of the QSAR model (Baguley, 2009). Internal validation exploits statistical methods in which different groups of chemicals are repetitively removed from the training set and biological activity of excluded compounds is predicted by the developed model. QSARINS employ Cross Validation (CV) parameters such as Q^2^ Leave One Out (Q^2^_LOO_), Q^2^ Leave Many Out (Q^2^_LMO_) and Root Mean Square Error (RMSE) for the verification of internal validation (Gramatica, 2007). Q^2^_LOO (leave one out)_ and Q^2^_LMO (leave many out)_ > 0.7 indicate models with high robustness and internal predictive ability (Douglas M. Hawkins et al., 2003). LMO cross validation is performed by eliminating nearly 20% of training set compounds in different cycles. Lower values of RMSE is also preferred for an acceptable QSAR model. Another validation approach employed by QSARINS includes Y-randomization, where the response values is shuffled randomly and models are generated. The parameters associated with Y-randomization include r^2^_Yscr_ and Q^2^_Yscr_. Lower values of these parameters indicate that models are not obtained by chance of correlation (Reads, 2017).

#### 2.4.2 External validation

The true predictive capacity of QSAR model is evaluated by carrying out external validation. External validation parameters obtained from QSARINS include r^2^_ext_, Q^2^F1, Q^2^F2 , Q^2^F3, CCC_ext_, RMSE_ext_, and r^2^m. Higher values of r^2^ (>0.6), Q^2^F1(>0.7), Q^2^F2(>0.7), Q^2^F3(>0.7), CCC_ext_ (0.85), r^2^m (>0.6) and lower values of RMSE_ext_ are acceptable for highly predictive QSAR model (Chirico and Gramatica, 2011). The true external predictive capacity of QSAR model is assessed by comparing the predicted and observed activities of an external set of compounds that were not used in QSAR model building. This is achieved by applying the QSAR model’s equation to one or more test data sets. The efficiency of QSAR model to predict activity of external dataset compounds is measured by analyzing the value of statistical parameter (r^2^).

### 2.5 Evaluation of Applicability Domain (AD)

AD is the chemical space defined by molecular descriptors and modeled responses. AD depicts the plot between leverage values and standard residuals. Leverage values are the diagonal elements of HAT matrix. AD helps in predicting uncertainty associated with a molecule by comparing its structural similarity to the compounds used to build the model (Netzeva et al., 2005). Compounds having unexpected biological activity and structural dissimilarity compared to other compounds in the dataset are considered outliers (Verma and Hansch, 2005).

### 2.6 Molecular docking studies

Molecular docking studies facilitate elucidation of the fundamental biochemical processes by enabling small molecule interactions with the active site of a protein. The active site of protein was identified by comparing the results obtained from computational tools such as Computed Atlas Surface Topography of proteins (CASTp), ScanProsite and existing literature references. CASTp identifies protein active site regions by considering the area and volume of the pocket by using a solvent accessible surface model and molecular surface model (Binkowski et al., 2003). ScanProsite detects functional and structural intradomain residues by utilizing context-dependent template annotations, which includes biological signatures such as expression patterns (de Castro et al., 2006). Molecular docking was performed by FlexX, which employs an incremental construction algorithm, and amalgamates a suitable model of the physico-chemical properties with effective methods for sampling the conformational space of the ligand. The algorithm aids exploring the binding properties of large numbers of flexible ligand conformers (Rarey et al., 1996). Molecular docking was proceeded with the definition of protein active site followed by providing preoptimized 3D coordinates of ligand. Identification of binding affinity and ligand efficiency was performed by the HYDE scoring function. HYDE scoring function uses hydrogen bond, torsion energies and desolvation terms of protein-ligand complexes for the prediction of binding affinity and ligand efficiency (Schneider et al., 2013).

### 2.7 Bioavailability assessment

The bioavailability assessment was carried out by StarDrop’s ADME module. Critical analysis of Absorption, Distribution, Metabolism, and Excretion provide significant insights to the complete metabolic profile of lead-like compounds. The substantial parameters characterizing bioavailability include logS (aqueous solubility), logP (partition coefficient), hERG pIC_50_ (cardiotoxicity), Human Intestinal Absorption (HIA), Plasma Protein Binding (PPB), cytochrome P450 affinities (CYP2C9 and CYP2D6). Aqueous solubility and partition coefficient are essential for proper absorption and distribution of compounds. Lower values of aqueous solubility and partition coefficient are not adequate for a good lead-like compound. hERG pIC_50_ is related to cardiotoxicity associated with the administration of drugs. hERG is a gene that encodes alpha subunit of the potassium channel, which plays a vital role in maintaining the electrical activity of the heart. Drug-induced inhibition of hERG contributes to the prolongation of QT interval, which could eventually lead to ventricular arrhythmias (Curran et al., 1995). Hence preclinical screening of lead molecules for hERG inhibition is highly essential. Positive absorption values and lower values of cytochrome affinities confirm good absorption and metabolic profile of compounds respectively. Binding of drugs with plasma proteins reduces their therapeutic efficacy by decreasing fractions reaching targeted cellular compartment. Plasma protein binding imparts a negative contribution to the efficient movement of the drug within cell membranes.

### 2.8 Identification of new derivatives

Identification of new derivatives was accomplished by REAL Space Navigator, tool which explores the largest chemical space comprising nearly 11 billion compounds using the building blocks from Enamine Ltd (Rarey and Dixon, 1998). The parent compound was considered as a reference molecule with pharmacophore constraints to generate derivatives. The Parent compound was selected from the experimental training dataset by setting a few checkpoints such as higher binding affinity, interactions with key amino acid residues of protein and enhanced bioavailability. Molecular docking and bioavailability assessments were carried out to identify binding interactions and effectiveness of derivative compounds compared to compounds in the experimental training dataset. Further, the potency of new derivatives to inhibit JAK2 was assessed by comparing molecular descriptor values and its physical significance obtained from validated QSAR model.

## 3. Results and discussion

### 3.1 QSAR model building

The splitting of compounds into training and prediction sets was based on similarity and clustering studies on 49 compounds. Compounds with ID S8000003, S8000004, S8000015, S8000020, S8000022, S8000023, S8000030, S8000032, S8000033, S8000039, S8000041, and S8000048 were classified as prediction set while rest of compounds were grouped as a training set. Validated QSAR model comprising 3 descriptors were obtained after the feature selection process. Compounds S8000016 and S8000024 were not included for QSAR model building due to their structural diversity with respect to other compounds present in the dataset.

Comprehensive information regarding dataset split is provided in (Supplementary Table S1).

### 3.2 Statistical and biological significance of QSAR model

The physical significance of descriptors and significant information regarding statistical parameters obtained from validated QSAR model is discussed in (**Table 1**).

**Table 1:**
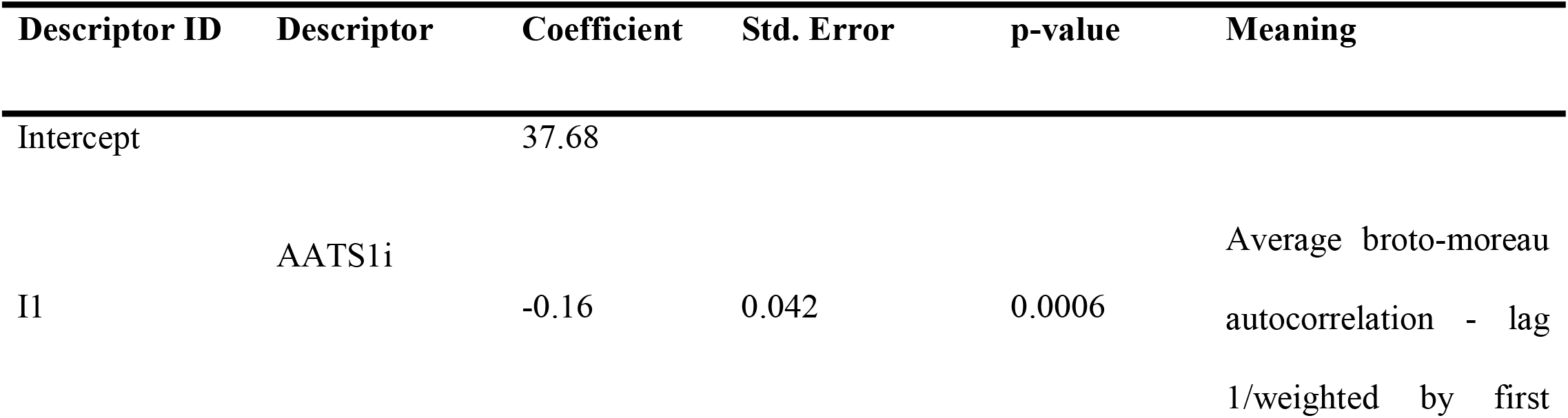
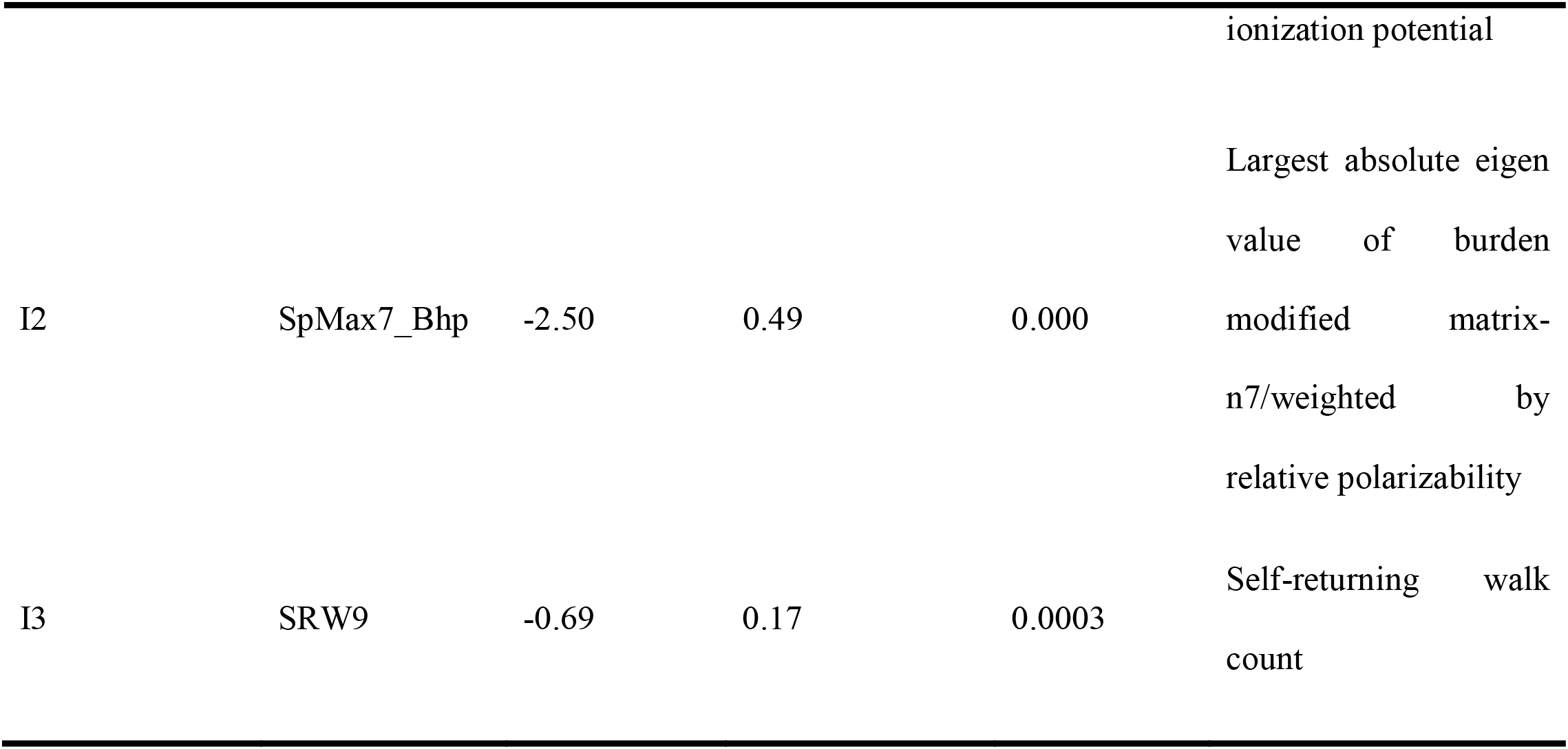
Physical significance of descriptors and validation parameters of QSAR model.

Parameters depicting fit of the model and internal validation

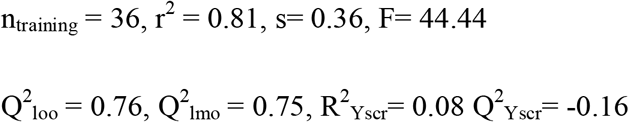

Predictivity statistics obtained by external validation

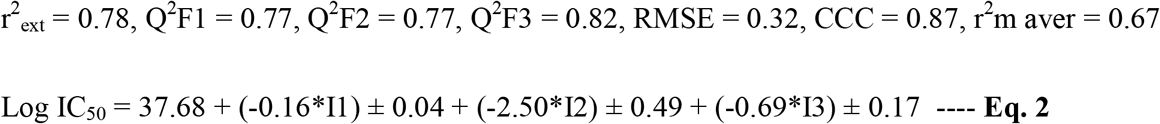

Permissible values of statistical parameters confirm the good fit and robustness of the QSAR model. Higher values of Q^2^-external and Concordance Correlation Coefficient (CCC) represents the predictive ability of the QSAR model. The inquisition of scatter plot and residual plot provide information regarding the statistical significance of QSAR model. Scatter plot and the residual plot are represented in (**Fig. 1**) And (**Fig. 2**) respectively.

**Fig. 1:**
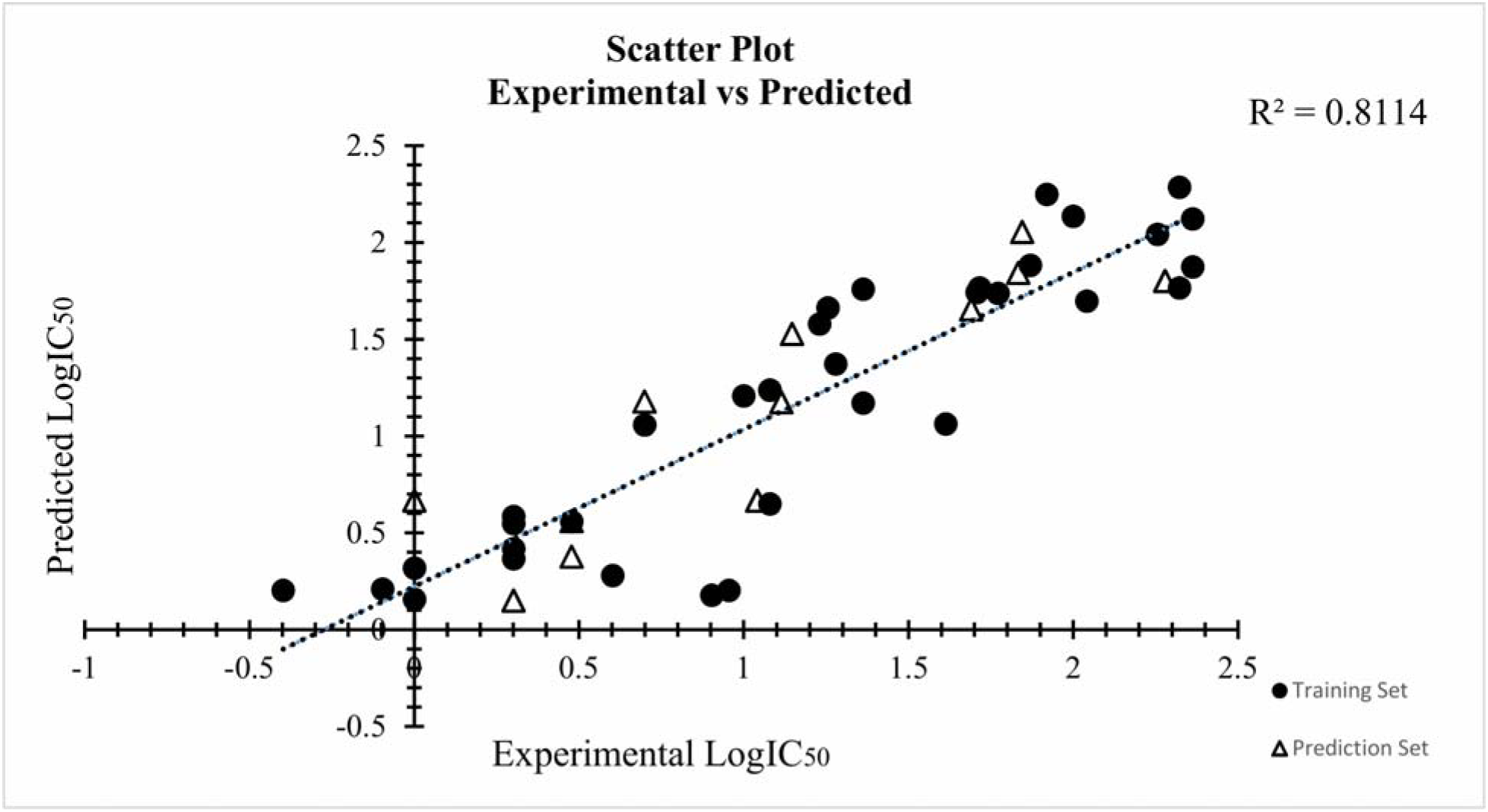
Depicts the correlation between experimental and predicted values for training set and prediction set compounds. Training set compounds are represented as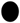 and prediction set compounds are depicted as Δ.

**Fig. 2:**
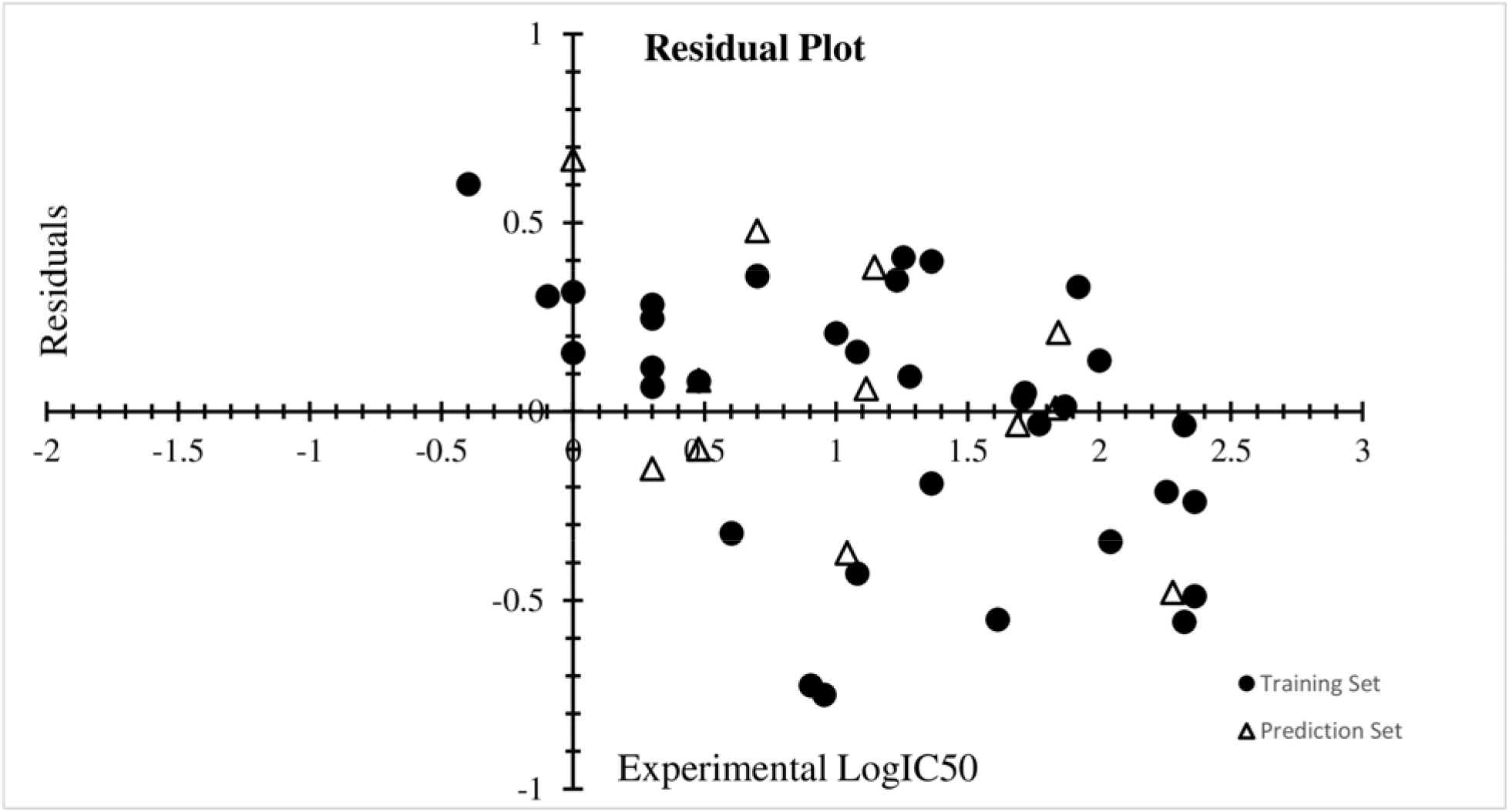
Illustrates correlation between experimental and residual values for training and prediction set compounds. The presence of randomly dispersed data points in a residual plot signifies the appropriateness of linear regression approach in modelling the data.

Key information regarding statistical parameters associated with QSAR model is provided in (Supplementary Table S2).

### 3.3 Physical Significance of descriptors

AATS1i, SpMax7_Bhp, and SRW9 were obtained as significant molecular descriptors. AATS1i (Average broto-moreau autocorrelation - lag 1/weighted by first ionization potential) belongs to the class of autocorrelation descriptors. Autocorrelation descriptors are topological descriptors which that that that encode structural and physico-chemical properties of a molecule (Sliwoski et al., 2016). This descriptor describes the distribution of a particular physicochemical property along its molecular structure. In this case, the property refers to ionization potential. Ionization potential refers to the ease of chemical bond formation by losing electrons from the valence shell of chemical compounds. SpMax7_Bhp (Largest absolute Eigen value of burden modified matrix-n7/weighted by relative polarizability) represent the class of burden modified Eigen value descriptors. These class of descriptors are defined as Eigen values of modified connectivity matrix known as Burden Matrix B. The diagonal elements of Burden Matrix B are set to account for different features of molecule such as electronegativity, polarizability etc. while off-diagonal elements represent bond orders (Todeschini et al., 2009). SpMax7_Bhp descriptor accounts for the polarizability of molecules used for QSAR model building. Polarizability accounts for the degree of ease by which electron cloud around an atom could be distorted. In general, negatively charged ions (anions) are highly polarizable and small-sized cations exhibit lower polarizability, but they possess the ability to polarize polarizable species such as anions (‘Polarizability - an overview | ScienceDirect Topics’, 2005). Polarizability plays a significant role in depicting nature of bonds formed between protein and ligand. The strength of protein-ligand interactions is always explained by the formation of a hydrogen bond, which results from a special type of dipole-dipole attraction between electronegative and electropositive atoms of ligand and protein. Hence, chemical compounds possessing polarizable or polarizing groups to enhance the degree of specific biochemical interactions with the target protein. Compounds in the dataset contain negatively charged atoms such as oxygen and nitrogen as well as smaller sized, positively charged hydrogen. This indicates the presence of both polarizable and polarizing groups in chemical compounds present in the dataset. Our docking studies demonstrated the role of these groups in making significant interactions with target protein JAK2. SRW9 (self-returning walk count descriptor) belongs to the class of walk and path count descriptor, which is obtained from the graph representation of the molecular structure. walk count of odd orders such as SRW9 accounts for the presence of odd-membered rings in chemical compounds. SRW9 descriptor also accounts for the size and nature of substituents attached to the odd membered ring (Todeschini and Consonni, 2000). All the compounds in the dataset have a 5 membered ring with NH-substituent. Hence, the QSAR model emphasizes on the significance of odd membered rings in eliciting JAK2 inhibition. Evidence from the literature suggest the presence of odd-numbered (specifically 5 membered rings) attached to heteroatoms in clinically proven JAK2 inhibitors such as ruxolitinib, baricitinib, pacritinib, lestaurtinib, and BMS-911543. Hence, our model also emphasizes the significance of odd-numbered rings in eliciting JAK2 inhibition.

Descriptor values for all compounds used for QSAR model building are provided in (Supplementary Table S3).

AD aids in the estimation of a model’s predictions with given reliability. No compounds were detected as outliers (HAT value:0.34). Hence, AD assessment revealed that all compounds in the dataset are influential in eliciting JAK2 inhibitory activity. William’s plot, which depicts relationship between leverage values and predicted values is shown in (**Fig. 3**).

**Fig. 3:**
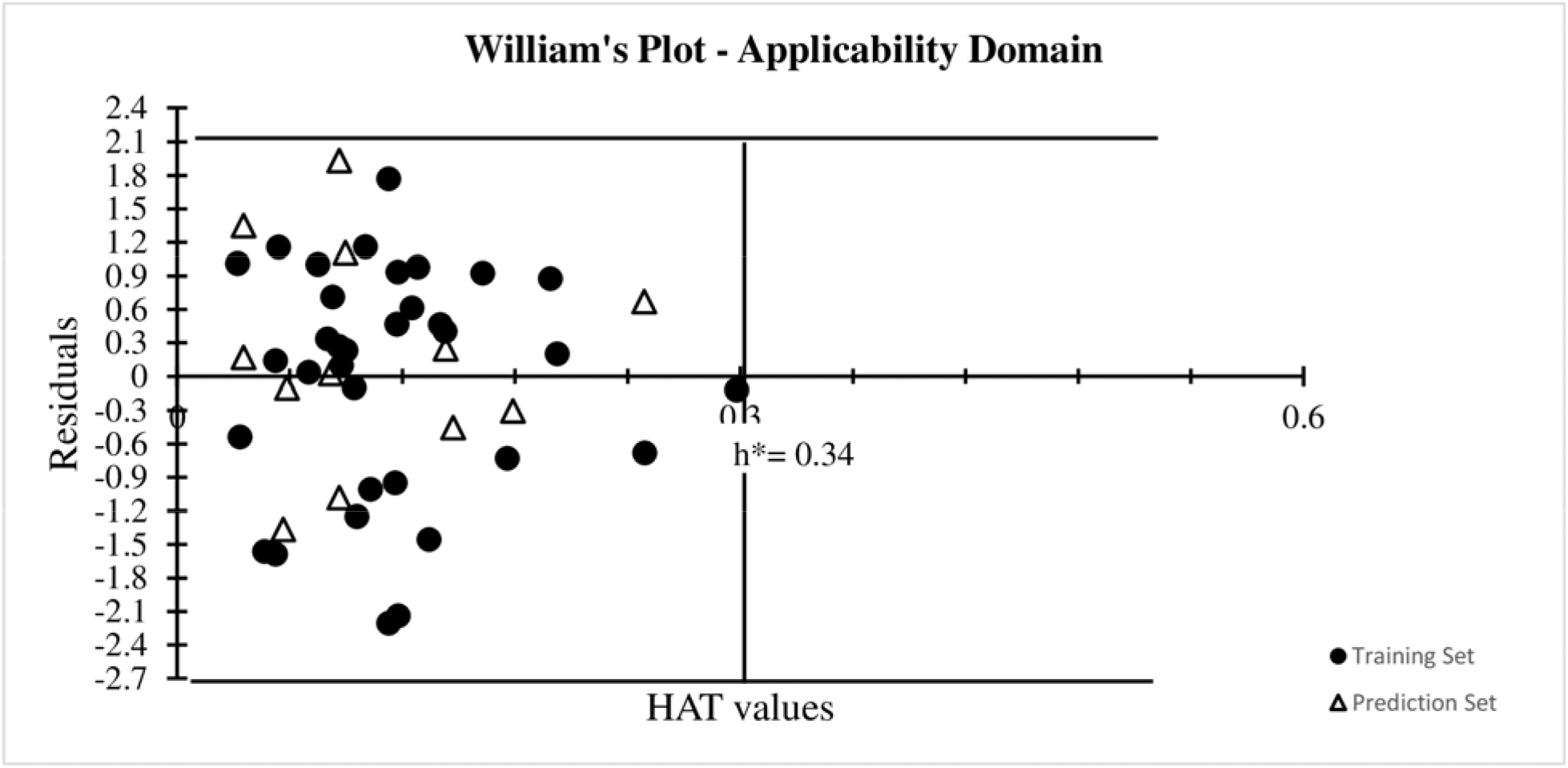
William’s plot depicting AD of the model. Leverage values are plotted on X-axis and residual values are plotted on Y-axis. Chemical compounds which don’t possess structural similarity with rest of compounds in the dataset are regarded as outliers.

QSAR validation plots are provided in Supplementary (Fig. S1-S4).

The true external predictive ability of QSAR model is evaluated by checking it’s ability to predict the activity of a set of compounds which were not used in QSAR model building. The set of 5 compounds was utilized for evaluating the external predictive capacity of QSAR model. The predictive ability of QSAR model was reflected from r^2^ value, which is 0.67. Molecular docking studies were also executed for these compounds to elucidate their mode of interactions with JAK2 protein. Docking studies revealed good binding affinity and significant interactions with JAK2 protein. The structures of compounds used for external validation and their descriptor values are provided in (Supplementary Table S4). Docking parameters associated with these compounds are provided in (Supplementary Table S5). Binding interactions (2D depiction)of compounds used for external validation with JAK2 protein are illustrated in Supplementary (Fig. S5).

### 3.4 Molecular docking studies

Molecular docking studies help in the elucidation of binding interactions between protein and ligands. The protein of interest was JAK2, (PDB ID:2B7A). Identification of active site residues was done from computational tools such as CASTp and ScanProsite. Active site residues identified from these two sources were compared with existing literature on JAK2 inhibitors. Met929, Leu855, Val863, Ala880, Val911, Leu983, Gly935, Tyr931, Glu930, Leu932, Asp939, Ser936, Arg980, Gly993, Asp994, Asn981, Asn859, Lys882, Phe860, and Asp976 were identified as active site residues. Out of these active site residues, Leu855, Val863, Ala880 and Val911 represented the class of hydrophobic residues occupied in the N-terminal lobe of JAK2 while Leu983 and Gly935 were present in C-terminal lobe. Met929 and Tyr931 were present in the hinge region of JAK2 (I.S. et al., 2006). Molecular docking studies using FlexX allowed to enable different stereo conformations of ligand molecule such as E/Z, R/S and pseudo R/S. 20,000 poses were generated per iteration including fragmentation and top 100 docking poses were analyzed.

Identification of JAK2 specific mutation in the onset and progression of MPNs lead towards the discovery of JAK2 specific inhibitors. Previously reported inhibitors targeted the ATP-binding pocket of JAK2 kinase domain (JH1). Studies conducted on the inhibitory activity of these compounds on JAK2 revealed the significance of interactions with hinge region residues and residues in the activation loop for eliciting a higher degree of potency (Burns et al., 2009),(Baskin et al., 2010). The interaction of inhibitors with Gly993 is highly preferred. Binding interaction with Gly993 provides a higher degree of specificity towards JAK2 since JAK3 possesses alanine residue in the same position (I.S. et al., 2006). Hence, screening of lead-like compounds was performed bymolecular docking studies and key amino acid residue interactions with the protein were analyzed. Compounds exhibiting optimal balance of dock score expected binding interactions as well as higher binding affinity with better torsion energies were considered for the next step of the analysis. Out of 49 compounds, 3 compounds S8000041, S8000042 and S8000032 exhibited binding affinity in the range of nanoMolar (nM) with JAK2 protein. Details regarding top 3 compounds is given (**Table 2**). Binding interactions of S8000041, S8000042 and S8000032 are provided in (**Fig. 4**, **5** **and** **6**) respectively.

**Table 2:** Depicts docking parameters associated with the binding of top 3 lead compounds with JAK2.

| Sl.no | Compound ID | Dock score | Binding Energy (KJ/mol) | Ligand efficiency | Binding affinity range | H-bond interactions (Amino acid residues) |
| --- | --- | --- | --- | --- | --- | --- |
| 1 | S8000041 | -27.17 | -34 | 0.28 | nM | Leu932, Tyr931, Met929, Gly993 |
| 2 | S8000042 | -26.9 | -35 | 0.28 | nM | Leu932, Glu930, Ser936 |
| 3 | S8000032 | -26.33 | -37 | 0.31 | nM | Leu932, Glu930 |

**Fig. 4:**
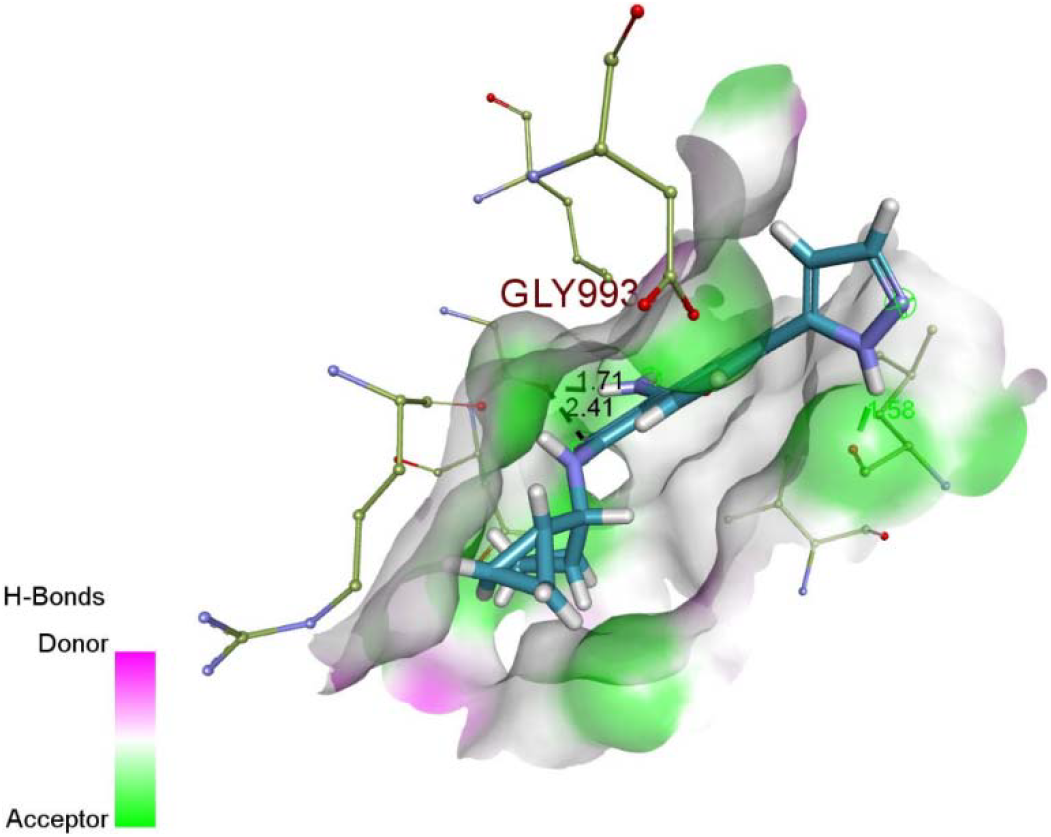
Binding mode of S8000041 with JAK2. S8000041 displayed hydrogen bond interactions with Leu932, Tyr931, Met929 and Gly993. Asn981, Ser936, Met929, Val863, Arg980, Asn981, Met929, Leu983, Ala880, Leu855, Tyr931 and Gly935 were found to exhibit hydrophobic interactions with S8000041.

**Fig. 5:**
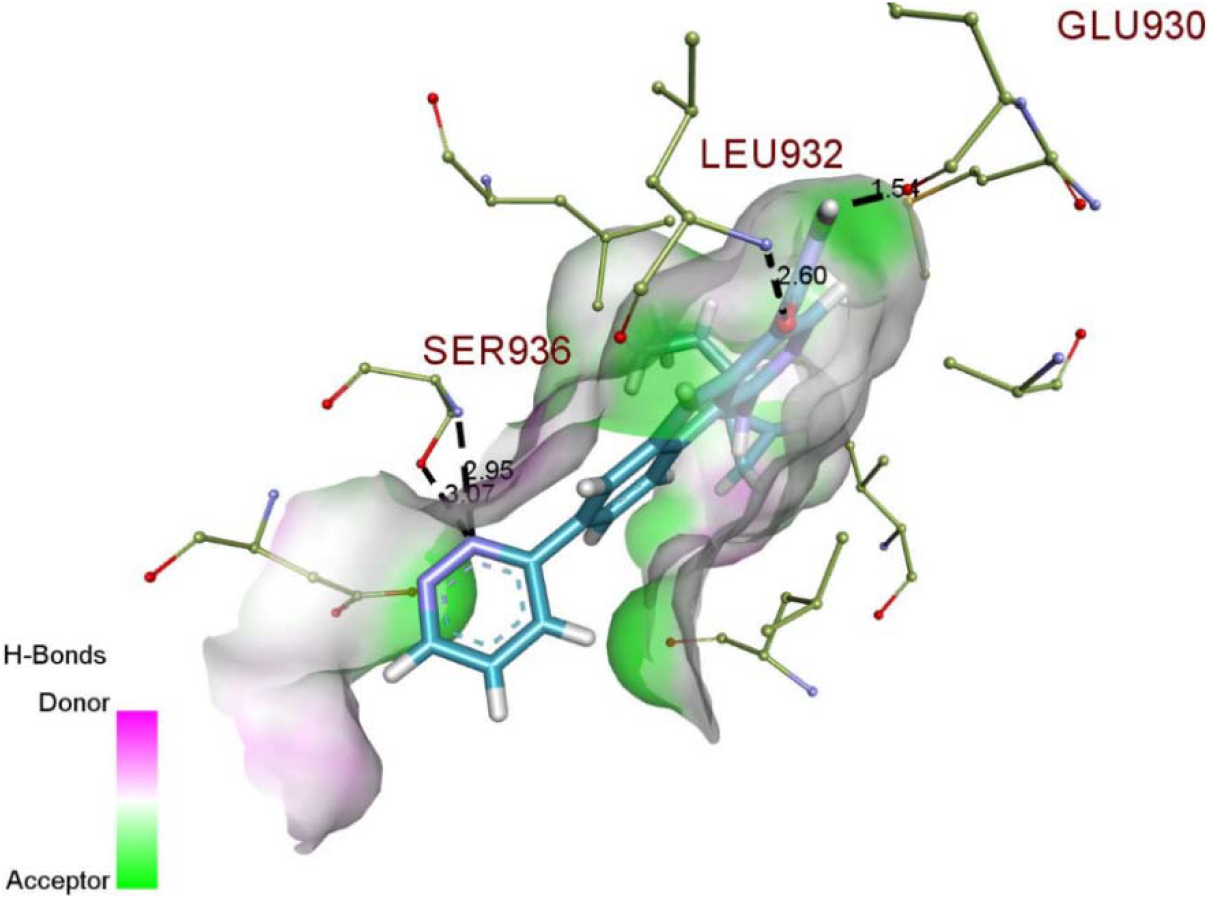
Depicts binding interactions of S8000042 with JAK2. Glu930, Leu932 and Ser936 exhibited hydrogen bond interactions with S8000042. Gly993, Val863, Asp994, Leu855, Leu983, Ser936 and Gly935 made hydrophobic interactions with S8000042.

**Fig. 6:**
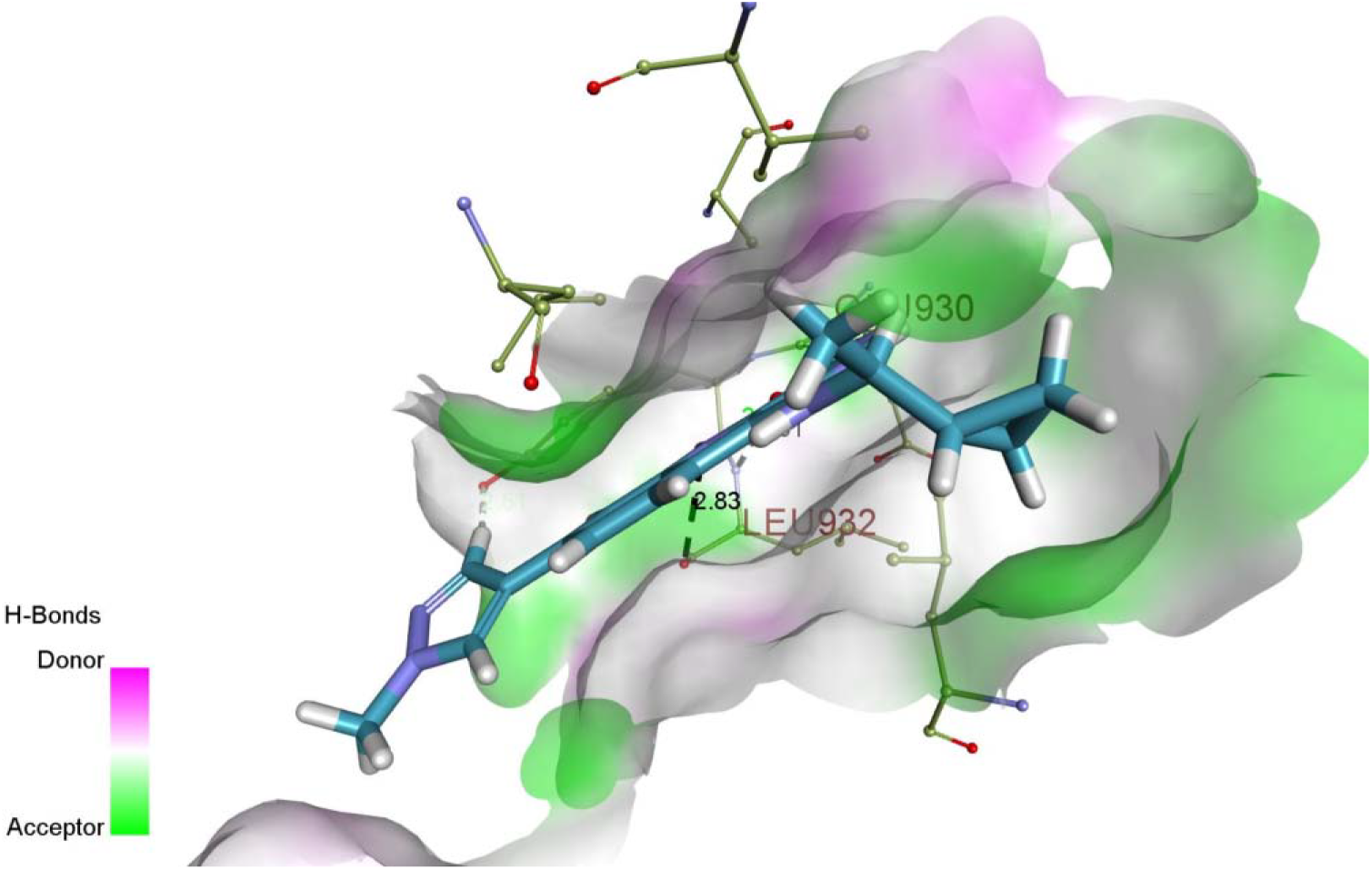
Binding interactions of S8000032. S8000032 revealed hydrogen bond interactions with Leu932 and Glu930, while Val863, Gly935, Leu855, Leu983, Tyr931 and Ala880 made hydrophobic interactions with JAK2 protein.

Compound S8000013 unveiled higher dock score of −33.01. But the binding energy, ligand efficiency as well as binding affinity range were found to be lesser for this compound. Hence, S8000013 was not selected as a lead like compound. Similarly, compounds having good dock scores which failed to satisfy other checkpoints such as good binding energy, ligand efficiency and binding affinity range were not considered as lead like compounds.

Out of top 3 compounds, S8000032 exhibited higher binding energy as well as ligand efficiency compared to other compounds. S8000032 established hydrogen bond interactions with active site residues such as Leu932 and Glu930. Leu932 is a hinge region residue that plays a significant role in eliciting the biological function of JAK2. But the hinge region is conserved across the JAK family, hence specificity towards JAK2 alone cannot be assured. But, S8000041 made significant interactions with hinge region residue as well as Gly993. The binding interaction with Gly993 confirmed specificity towards JAK2 since alanine is present in the same position for other members of JAK family. Met929 is known as “gatekeeper residue”. Previously reported inhibitor CMP6 developed by Merck Research Laboratories exhibited interactions with the gatekeeper residue. The gate keeper residue plays a vital role in maintaining the shape and size of the binding pocket. Met 929 constricts the active site and enhances shape complementarity for facilitating interactions with specific ligand groups (I.S. et al., 2006). Hence, compound S8000041 displayed significant binding affinity and biological significance for eliciting JAK2 inhibition.

Dock score, binding energy, ligand efficiency and binding affinity range for all compounds present in dataset are provided in (Supplementary Table S6).

Binding interactions (2D depiction) of all 49 compounds are given in Supplementary (Fig. S6).

### 3.5 Bioavailability assessment

StarDrop performs bioavailability prediction based on the inbuilt QSAR models for each parameterdepicting bioavailability (Shen et al., 2010). The threshold values for bioavailability parameters are as follows: solubility (logS) > 1, partition coefficient (logP) < 3.5, affinity towards cytochrome p450 isoforms - low/medium, Human Intestinal Absorption (HIA +ve), 2C9 pKi < 6, hERG liability - <5 and plasma protein binding – low. All three compounds exhibited exceptional values for physicochemical features representing bioavailability such as solubility, partition coefficient, affinity towards cytochrome isoform 2C9 and hERG liability. But affinity towards cytochrome isoform 2D6 and plasma protein binding were found to be very high. Higher affinity towards cytochrome p450 and plasma protein hampers the efficacy of the drug to elicit targeted biological function and proper distribution in cellular compartments. Hence, there is an urgent need to identify lead compounds with better potency and bioavailability. Out of the three lead compounds identified, solubility was found to be higher for S8000032 but S8000041 showed excellent values for other physicochemical features predicting bioavailability closer to their threshold values. Docking studies proved that S8000041 displayed significant biological interactions with JAK2 protein. Hence this compound was used as a parent compound for the identification of derivatives. Bioavailability parameters for the top 3 lead compounds, which exhibited good binding affinity and significant interactions with target protein is given in (**Table 3**).

**Table 3:** Provides significant information regarding bioavailability parameters for the top 3 lead compounds.

| Sl.no | Compound ID | logS | logP | 2C9 pKi | hERG pIC50 | HIA | 2D6 affinity | PPB90 category |
| --- | --- | --- | --- | --- | --- | --- | --- | --- |
| 1 | S8000041 | 1.416 | 3.018 | 5.1 | 4.35 | + | High | High |
| 2 | S8000042 | 1.337 | 3.88 | 5.3 | 4.9 | + | Very high | High |
| 3 | S8000032 | 1.298 | 3.4 | 5.6 | 4.6 | + | High | High |

Bioavailability assessment results for all the compounds in the dataset is provided in (Supplementary Table S7).

### 3.6 Identification of new derivatives

Derivatives for S8000041 compound were identified from REAL Space Navigator, which is the world’s largest ultra-fast searchable chemical space. This database is built based on 121 enamine synthesis protocols and in-stock building blocks. 1000 derivative compounds were predicted for S8000041 by fixing tanimoto coefficient at 0.8. The new derivatives ensure synthetic feasibility and accessibility checks to ensure the compounds are synthesizable.

Vital information regarding the structure and physicochemical properties associated with derivative compounds of S8000041 is provided in (Supplementary Table S8).

### 3.7 Molecular docking studies

Molecular docking studies were executed to identify the binding efficacy of 1000 derivative compounds with JAK2 protein (PDB ID: 2B7A). The definition of active sites was followed by molecular docking and binding affinity prediction (HYDE assessment). Details regarding docking scores, binding energy values, ligand efficiency and binding affinity range are provided in (**Table 4**). Intermediate compounds represent building blocks (ID referred from www.enamine.net) is used for entire compound by Enamine.

**Table 4:** Provides significant insights into binding mode of top 10 derivative compounds with JAK2 protein.

| Compound ID | Building blocks (for synthesis) | Dock score | Binding Energy (kJ/mol) | Ligand efficiency | Binding affinity range |
| --- | --- | --- | --- | --- | --- |
| D1 | S2708-1034522-14187958 | -41.02 | -38 | 0.35 | nM |
| D2 | S271948-9942056-9919210 | -39.73 | -24 | 0.21 | μM-nM |
| D3 | S271949-14780076-12857078 | -39.68 | -19 | 0.17 | μM-nM |
| D4 | S2708-7026222-11967138 | -39.65 | -26 | 0.23 | μM-nM |
| D5 | S2430-15146132-9964506 | -39.27 | -27 | 0.22 | $\mu\text{M}$-nM |
| D6 | S271948-9948308-9919210 | -39.24 | -11 | 0.09 | mM-$\mu\text{M}$ |
| D7 | S271949-10832104-12857078 | -38.93 | -14 | 0.13 | mM-$\mu\text{M}$ |
| D8 | S2708-1001846-14187958 | -38.83 | -25 | 0.23 | $\mu\text{M}$-nM |
| D9 | S2708-12122826-14187958 | -38.56 | -23 | 0.2 | $\mu\text{M}$-nM |
| D10 | S272212-13250008-9425858 | -34.97 | -17 | 0.14 | mM-$\mu\text{M}$ |

From the set of 1000 compounds, top 10 compounds possessing higher dock scores were screened initially. These compounds were subjected to HYDE assessment for analyzing binding energy, ligand efficiency and binding affinity range. D1 established higher dock score, binding energy as well as ligand efficiency. The binding affinity range of this compound was predicted to be in Nano molar range. H-bond interaction with Gly993 accounted for the specificity of D1 towards JAK2. In addition to this, literature on JAK2 inhibitors reported that interaction with Asp939 is also responsible for maintaining a higher degree of specificity towards JAK2 (Zhao et al., 2016). Similarly compounds D2, D3, D4, D5, D6, D7, D8, D9 and D10 exhibited interactions with Asp939, which explained the specificity of these compounds towards JAK2. Even though D10 showed comparatively lower dock score, binding energy, ligand efficiency and binding affinity range it made significant interactions with Gly993 and Asp939. (**Fig. 7** **and** **8**) represent binding interactions of derivative compounds D2 and D5 respectively.

**Fig. 7:**
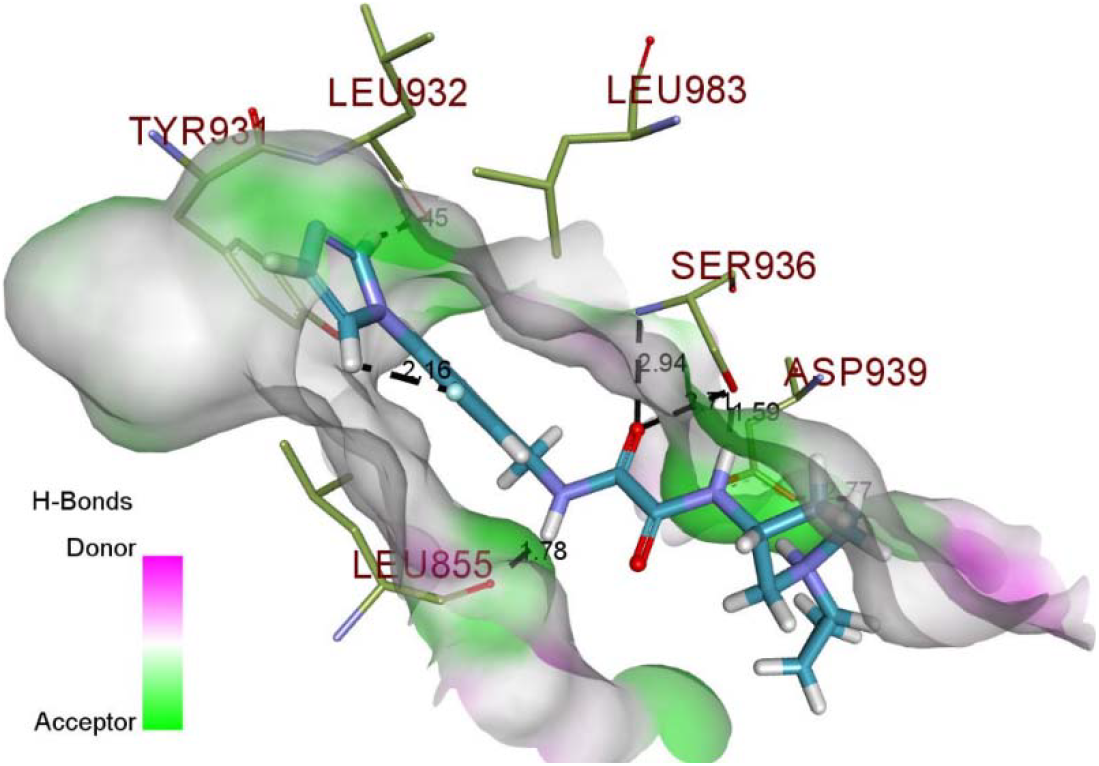
Depicts binding interactions of derivative compound D2 with JAK2 protein. D2 formed H-bond interactions with Asp939, Leu855, Leu932 and Ser936. Compound formed hydrophobic interactions with Gly935, Leu855, Tyr931, Val863, Leu983, Ser936 and Arg980.

**Fig. 8:**
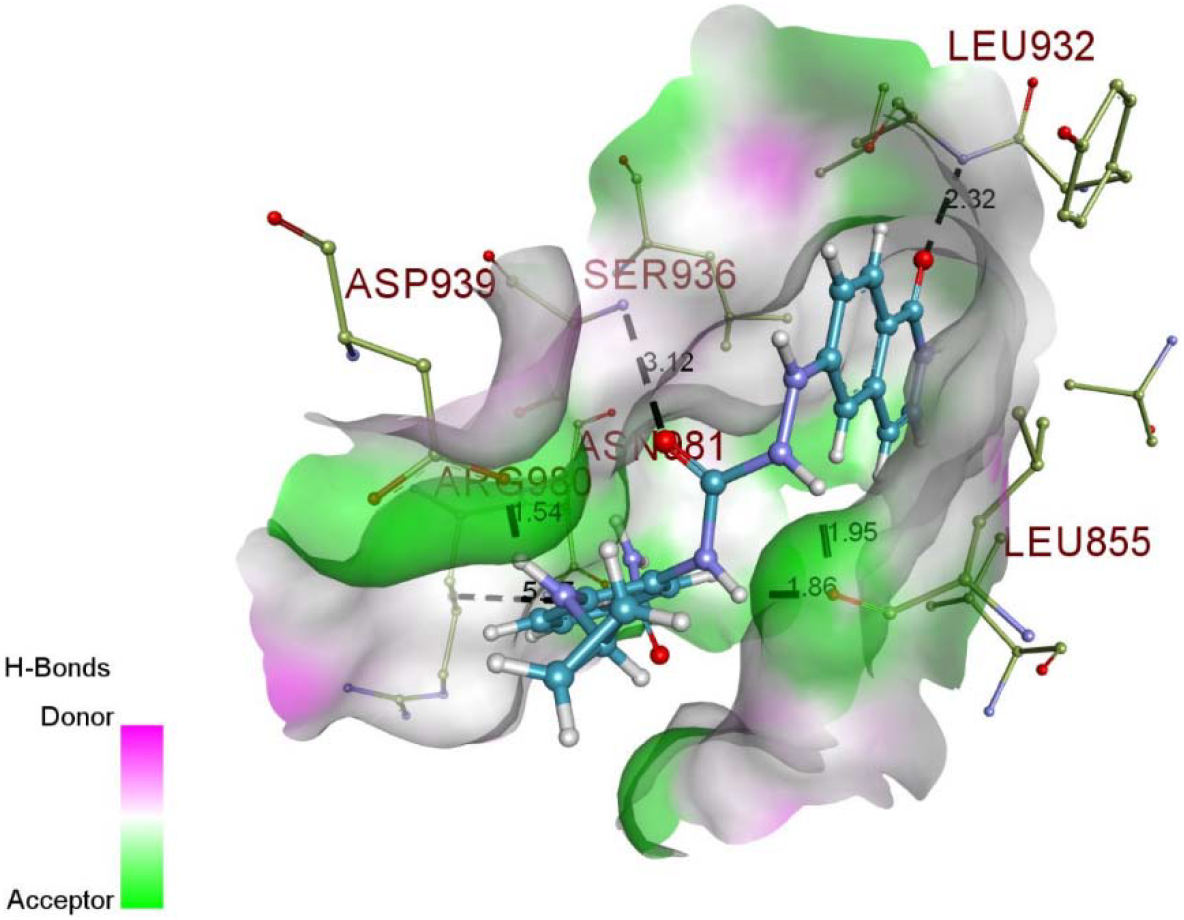
Derivative compound D5 made H-bond interactions with Asp939, Leu855, Leu932, Arg980, Asn981 and Ser936, while hydrophobic interactions were made with Gly935, Tyr931, Arg980, Ser936, Leu984, Leu863, Leu983 and Leu855.

### 3.8 Bioavailability assessment for top 10 derivatives

Out of top 10 compounds, D2, D3, D5, D6, and D7 showed outstanding bioavailability since the predictions were within the threshold values (logS > 1, logP < 3.5, affinity towards cytochrome P450 isoforms - low/medium, Human Intestinal Absorption (HIA +ve), 2C9 pKi < 6, hERG liability - <5 and plasma protein binding – low). Compound D1, showed good binding affinity, significant interactions with the target protein and appreciable bioavailability (good solubility, lipophilicity, human intestinal absorption, and lower plasma protein binding profiles). But affinity towards cytochrome P450 isoform 2D6 was significantly higher and there was a marginal increase in hERG liability. StarDrop’s glowing molecule module predicted the structural features of D1, responsible for the enhanced affinity towards cytochrome p450. The presence of trimethyl groups, nitrogen atoms, carbonyl groups, and methylene groups are primarily responsible for the increased affinity towards cytochrome p450. The marginal increase in the value of hERG pIC_50_ was attributed to the presence of trimethyl groups, nitrogen atoms, carbonyl group, benzene ring and piperidine group. Similarly, compound D10 which showed interactions with Gly993 demonstrated marginal increase in hERG liability. hERG liability was associated with the presence of piperidine group, nitrogen atom, methylene groups and 4-bromo-indole groups. (**Fig. 9)** depicts the structural features of derivative compounds contributing for bioavailability parameters. The structural features of D1 responsible for upregulated bioavailability parameters such as cytochrome p450 and hERG liability are provided in (**Fig. 9a** **and** **9b)** respectively. (**Fig. 9c)** depicts structural features of D10 responsible for the increased values of hERG liability.

**Fig. 9:**
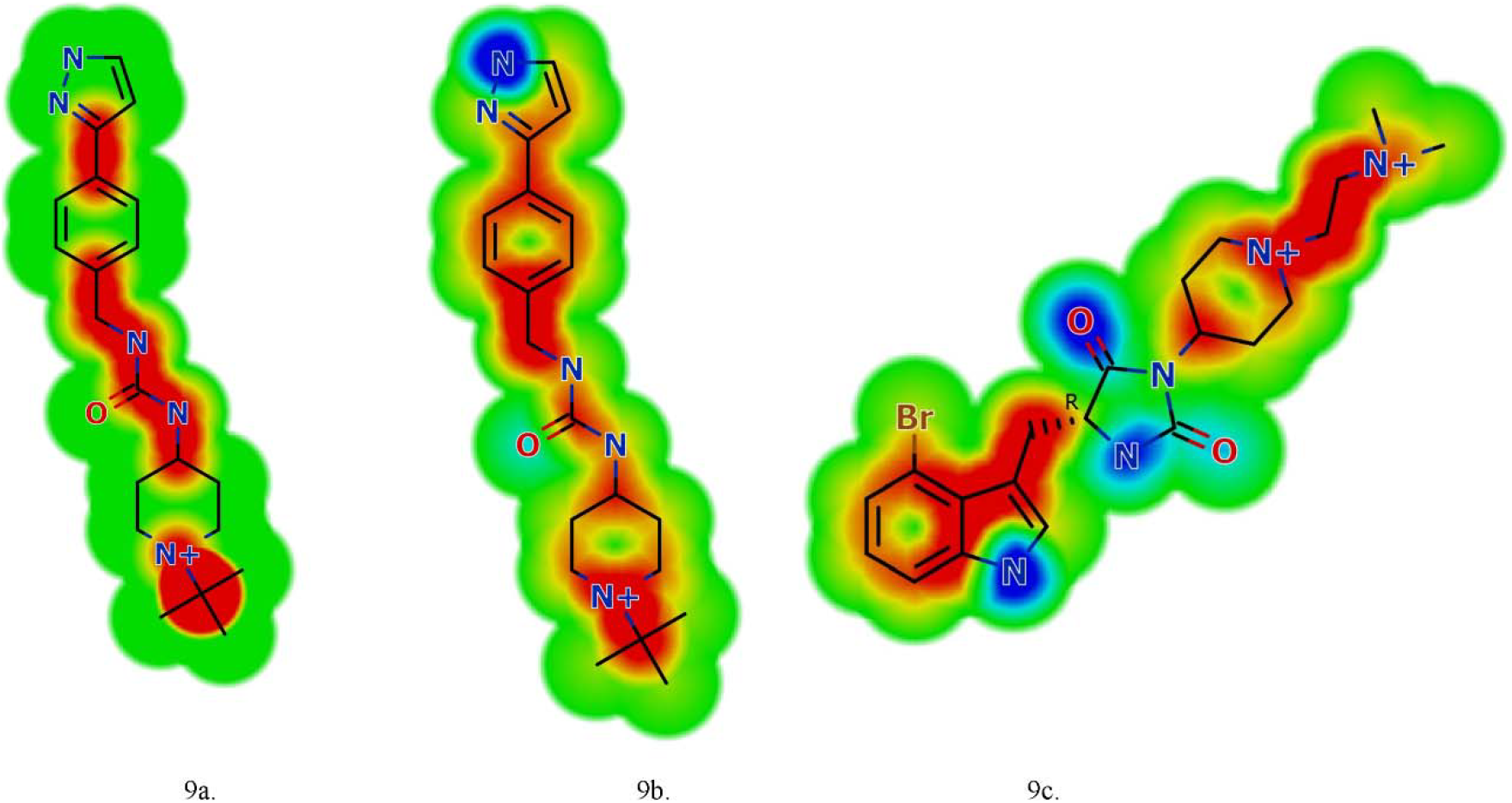
Structural features of derivative compounds D1 and D10 contributing for bioavailability parameters. **(Fig. 9a)** Depicts structural features of derivative compound D1 contributing for bioavailability parameter cytochrome p450 isoform 2D6. (**Fig. 9b)** Depicts structural features of derivative compound D1 contributing for bioavailability parameter hERG liability. (**Fig. 9c)** Depicts structural features of derivative compound D10 contributing for bioavailability parameter hERG liability.

But all other bioavailability parameters were predicted within the threshold range. But compounds D2 and D5 showed substantial binding affinity as well as bioavailability among the top 10 derivative compounds. Hence, we propose D2 and D5 as potential JAK2 inhibitors by considering multiple parameters such as binding affinity, binding interactions as well a bioavailability. (**Table 5**). Depicts physicochemical properties predicting bioavailability for top 10 derivative compounds.

**Table 5:**
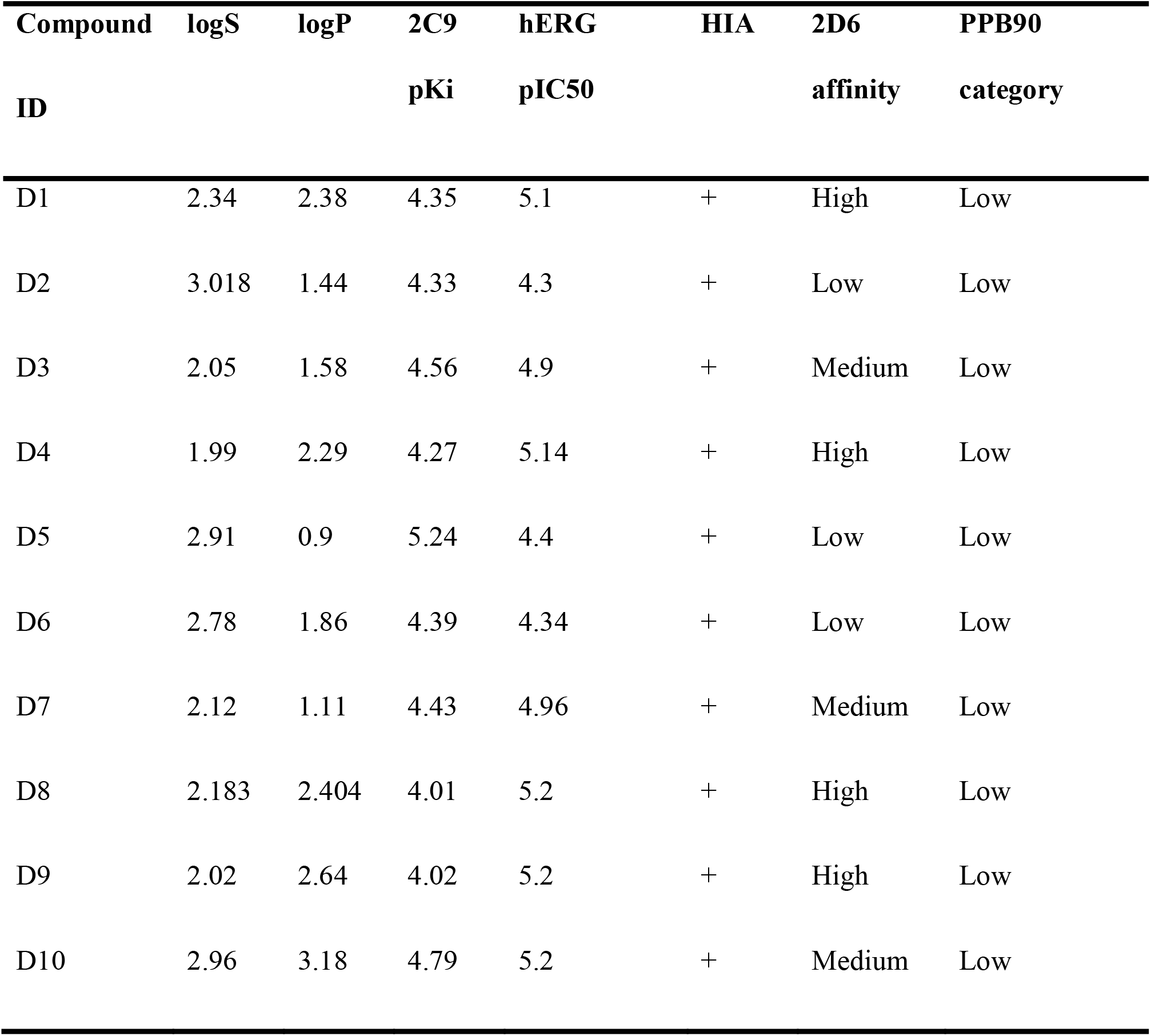
Details regarding ADME prediction for top 10 derivative compounds.

### 3.9 Activity prediction of derivative compounds using QSAR model

Validated QSAR model reported the significance of 3 descriptors in eliciting JAK2 inhibition. These descriptors were calculated for the top 10 derivative compounds. Comparative analysis of these descriptor values with that obtained from a validated QSAR model suggested similarity. The similarity of descriptor values of derivative compounds with respect to compounds used in QSAR model building propose similarity in biological function as well.

Descriptor values for the top 10 derivative compounds are provided in (Supplementary Table S9).

## 4. Conclusion

This study aimed to identify novel JAK2 inhibitors possessing improved biological activity as well as bioavailability. We integrated ligand and structural based studies to propose novel inhibitors against JAK2. Initially, QSAR model was developed using chemical compounds reported for JAK2 inhibitory activity. Validated QSAR model proposed the significance of odd-numbered rings in chemical compounds for eliciting JAK2 inhibition. This observation was confirmed by comparing the structural features of clinically proven JAK2 inhibitors. Further, the descriptor present in validated QSAR model revealed the significance of polarizability in establishing stronger hydrogen bonds with specific amino-acid residues present in the JAK2 active site. Hit to lead optimization strategies included molecular docking and bioavailability assessments. The interaction of compounds with specific amino acid residues confered their potency and specificity for eliciting JAK2 inhibition. Three compounds were identified as potential lead compounds by considering the above mentioned criteria. S8000041 was found as the most promising lead molecule. But all 3 compounds were found to have higher metabolic profiles which lead to faster degradation of these compounds before eliciting its therapeutic action. Hence derivative compounds for S8000041 were identified from the Enamine database.

Hit to lead optimization strategies were performed similarly as done for parent compounds and the top 10 derivative compounds were identified. Consideration of multiple parameters such as binding affinity and bioavailability resulted in the identification of 2 lead molecules (D2 and D5). These compounds exhibited interactions with Asp939, which is found as a significant interaction in eliciting potency and specificity towards JAK2 inhibition. The structural features of chemical compounds play significant role in eliciting targeted biological activity. Hence, descriptors obtained from validated QSAR were compared with that of derivative compounds to identify their structural similarity. Comparative analysis of descriptor values confirmed structural similarity of derivative compounds with the compounds (reported JAK2 inhibitors) employed for QSAR model building. The similarity in descriptor values denotes the significance of odd-numbered rings and presence of polarizing groups in making specific hydrogen bond interactions with JAK2 protein. Hence, our studies propose two derivative compounds (D2 and D5) as potential JAK2 inhibitors. Further, *invitro* and *invivo* studies has to be conducted for assuring the potency of these compounds as JAK2 inhibitors.

## Supporting information

SI - Dataset Docking QSAR Data

## Author contributions

GGP aided in the conceptualization of the study and provided resources. AUP carried-out experiments, analysis and wrote manuscript. SLS analysed and validated data.

## Acknowledgements

The authors thank Optibrium Ltd, UK, and BioSolveIT GmbH, Germany for providing licenses for StarDrop, FlexX, RealSpaceNavigator software tools. Authors extend their thanks to Prof. Paula Gramatica and Prof. Alan R. Katritzky for permitting us to use QSARINS and CodessaPRO software for this work. Author Ambili thank Vellore Institute of Technology for immense financial support for this research work and thank Arushi Mishra for language improvement.

## Supplementary Material

