## Supplementary material for "Integrated Ligand and Structure based approaches towards developing novel Janus Kinase 2 inhibitors for the treatment of myeloproliferative neoplasms": SI - Dataset Docking QSAR Data

We have applied integrated ligand and structure-based approaches for developing novel JAK2 inhibitors. Ligand based study (QSAR modelling) was employed to elucidate the physicochemical properties of chemical compounds responsible for eliciting JAK2 inhibition. Further molecular docking and bioavailability assessments were performed to identify most promising lead compounds. Derivative compounds which possess excellent potency and bioavailability than the existing compounds reported for JAK2 inhibition were identified.

**Table S1**: Depicts details regarding experimental and predicted IC_50_ values for chemical compounds classified as training and prediction set in QSAR modelling.

| **ID** | **Structure** | **Exp-log IC_50_** | **pIC_50_** |
| --- | --- | --- | --- |
| S8000001^T^ | C1=C(C2=C(C(=N1)NC)C3=C([NH]2)C=CC(=C3)F)C(=O)N | 2.322 | 2.2862 |
| S8000002^T^ | C1=C(C2=C(C(=N1)NCC)C3=C([NH]2)C=CC(=C3)F)C(=O)N | 2.362 | 2.1232 |
| S8000003^P^ | C1=C(C2=C(C(=N1)NCCC)C3=C([NH]2)C=CC(=C3)F)C(=O)N | 1.833 | 1.842 |
| S8000004^P^ | C1=C(C2=C(C(=N1)NCCCC)C3=C([NH]2)C=CC(=C3)F)C(=O)N | 1.69 | 1.6548 |
| S8000005^T^ | C1=C(C2=C(C(=N1)NC(C)C)C3=C([NH]2)C=CC(=C3)F)C(=O)N | 2 | 2.1352 |
| S8000006^T^ | C1=C(C2=C(C(=N1)NCC(C)C)C3=C([NH]2)C=CC(=C3)F)C(=O)N | 1.869 | 1.8829 |
| S8000007^T^ | C1=C(C2=C(C(=N1)N[C@H](C(C)(C)C)C)C3=C([NH]2)C=CC(=C3)F)C(=O)N | 1.716 | 1.7657 |
| S8000008^T^ | C1=C(C2=C(C(=N1)N[C@@H](C(C)(C)C)C)C3=C([NH]2)C=CC(=C3)F)C(=O)N | 2.322 | 1.7657 |
| S8000009^T^ | C1=C(C3=C(C(=N1)NC2CCC2)C4=C([NH]3)C=CC(=C4)F)C(=O)N | 1.919 | 2.2497 |
| S8000010^T^ | C1=C(C3=C(C(=N1)NC2CCCC2)C4=C([NH]3)C=CC(=C4)F)C(=O)N | 1.708 | 1.7427 |
| S8000011^T^ | C1=C(C3=C(C(=N1)NC2CCCCC2)C4=C([NH]3)C=CC(=C4)F)C(=O)N | 1.23 | 1.5787 |
| S8000012^T^ | C1=C(C3=C(C(=N1)NC2CCCCCC2)C4=C([NH]3)C=CC(=C4)F)C(=O)N | 1.279 | 1.3721 |
| S8000013^T^ | C1=C(C3=C(C(=N1)NC2CCOCC2)C4=C([NH]3)C=CC(=C4)F)C(=O)N | 1.255 | 1.6629 |
| S8000014^T^ | C1=C(C3=C(C(=N1)N2CCCC2)C4=C([NH]3)C=CC(=C4)F)C(=O)N | 2.362 | 1.8737 |
| S8000015^P^ | C1=C(C3=C(C(=N1)N2CCCCC2)C4=C([NH]3)C=CC(=C4)F)C(=O)N | 2.279 | 1.8005 |
| S8000017^T^ | C1=NC(=C2C(=C1C(=O)N)[NH]C3=C2C=CC=C3)NC4CCCCC4 | 1.771 | 1.7377 |
| S8000018^T^ | C1=NC(=C2C(=C1C(=O)N)[NH]C3=C2C=CC(=C3)Cl)NC4CCCCC4 | 1.362 | 1.7601 |
| S8000019^T^ | C1=NC(=C2C(=C1C(=O)N)[NH]C3=C2C(=CC=C3)Cl)NC4CCCCC4 | 2.041 | 1.697 |
| S8000020^P^ | C1=NC(=C2C(=C1C(=O)N)[NH]C3=C2C=CC(=C3)C4=CC=CC=C4)NC5CCCCC5 | 1.845 | 2.0546 |
| S8000021^T^ | C1=NC(=C2C(=C1C(=O)N)[NH]C3=C2C=C(C=C3)C4=CC=CC=C4)NC5CCCCC5 | 2.255 | 2.0421 |
| S8000022^P^ | C1=NC(=C2C(=C1C(=O)N)[NH]C3=C2C=CC(=C3)C4=CC=N[N]4C)NC5CCCCC5 | 0.699 | 1.1771 |
| S8000023^P^ | C1=NC(=C2C(=C1C(=O)N)[NH]C3=C2C=C(C=C3)C4=C[N](N=C4)C)NC5CCCCC5 | 1.114 | 1.1746 |
| S8000025^T^ | C1=NC(=C2C(=C1C(=O)N)[NH]C3=C2C=CC(=C3)C4=CC=N[NH]4)NC5CCCCC5 | 1.362 | 1.1717 |
| S8000026^T^ | C1=NC(=C2C(=C1C(=O)N)[NH]C3=C2C=CC(=C3)C4=C[N](N=C4)C)N[C@H](C(C)(C)C)C | 0.699 | 1.0585 |
| S8000027^T^ | C1=NC(=C2C(=C1C(=O)N)[NH]C3=C2C=CC(=C3)C4=C[N](N=C4)CC(C)C)N[C@H](C(C)(C)C)C | 1.613 | 1.0631 |
| S8000028^T^ | C1=NC(=C2C(=C1C(=O)N)[NH]C3=C2C=CC(=C3)C4=CN=C(N=C4)N)N[C@H](C(C)(C)C)C | 1.079 | 1.2379 |
| S8000029^T^ | C1=NC(=C2C(=C1C(=O)N)[NH]C3=C2C=CC(=C3)C4=CC=NC(=N4)N)N[C@H](C(C)(C)C)C | 1.079 | 1.2379 |
| S8000030^P^ | C1=NC(=C2C(=C1C(=O)N)[NH]C3=C2C=CC(=C3)C4=CC=C(N=N4)Cl)N[C@H](C(C)(C)C)C | 1.146 | 1.528 |
| S8000031^T^ | C1=NC(=C2C(=C1C(=O)N)[NH]C3=C2C=CC(=C3)C4=CC=C(N=N4)N)N[C@H](C(C)(C)C)C | 1 | 1.208 |
| S8000032^P^ | C1=C(C3=C(C(=N1)N[C@H](C2CC2)C)C4=C([NH]3)C=C(C=C4)C5=C[N](N=C5)C)C(=O)N | 0 | 0.6672 |
| S8000033^P^ | C1=C(C3=C(C(=N1)N[C@@H](C2CC2)C)C4=C([NH]3)C=C(C=C4)C5=C[N](N=C5)C)C(=O)N | 1.041 | 0.6672 |
| S8000034^T^ | C1=C(C3=C(C(=N1)N[C@H](C2CC2)CC)C4=C([NH]3)C=C(C=C4)C5=C[N](N=C5)C)C(=O)N | 0.301 | 0.548 |
| S8000035^T^ | C1=C(C3=C(C(=N1)N[C@H](C2CC2)C(C)C)C4=C([NH]3)C=C(C=C4)C5=C[N](N=C5)C)C(=O)N | 0.477 | 0.558 |
| S8000036^T^ | C1=C(C4=C(C(=N1)NC(C2CC2)C3CC3)C5=C([NH]4)C=C(C=C5)C6=C[N](N=C6)C)C(=O)N | 0.301 | 0.3669 |
| S8000037^T^ | C1=C(C3=C(C(=N1)N[C@H](C2CC2)C(F)(F)F)C4=C([NH]3)C=C(C=C4)C5=C[N](N=C5)C)C(=O)N | -0.398 | 0.2041 |
| S8000038^T^ | C1=C(C3=C(C(=N1)N[C@@H](C2CC2)C(F)(F)F)C4=C([NH]3)C=C(C=C4)C5=C[N](N=C5)C)C(=O)N | 0.954 | 0.2041 |
| S8000039^P^ | C1=C(C2=C(C(=N1)N[C@H](C(C)C)C(F)(F)F)C3=C([NH]2)C=C(C=C3)C4=C[N](N=C4)C)C(=O)N | 0.477 | 0.5594 |
| S8000040^T^ | C1=C(C3=C(C(=N1)N[C@H](C2CCC2)C(F)(F)F)C4=C([NH]3)C=C(C=C4)C5=C[N](N=C5)C)C(=O)N | 1.079 | 0.6505 |
| S8000041^P^ | C1=C(C4=C(C(=N1)NC(C2CC2)C3CC3)C5=C([NH]4)C=C(C=C5)C6=CC=N[NH]6)C(=O)N | 0.477 | 0.3772 |
| S8000042^T^ | C1=C(C4=C(C(=N1)NC(C2CC2)C3CC3)C5=C([NH]4)C=C(C=C5)C6=CC=CN=N6)C(=O)N | 0.301 | 0.5851 |
| S8000043^T^ | C1=C(C4=C(C(=N1)NC(C2CC2)C3CC3)C5=C([NH]4)C=C(C=C5)C6=CN=C(C=N6)N)C(=O)N | 0.602 | 0.2802 |
| S8000044^T^ | C1=C(C4=C(C(=N1)NC(C2CC2)C3CC3)C5=C([NH]4)C=C(C=C5)C6=CN=C(N=C6)N)C(=O)N | 0 | 0.3178 |
| S8000045^T^ | C1=C(C3=C(C(=N1)N[C@@H](C(F)(F)F)C2CC2)C4=C([NH]3)C=C(C=C4)C5=C[NH]N=C5)C(=O)N | 0 | 0.1562 |
| S8000046^T^ | C1=C(C3=C(C(=N1)N[C@@H](C(F)(F)F)C2CC2)C4=C([NH]3)C=C(C=C4)C5=CN=C(C=C5)N)C(=O)N | 0.301 | 0.4182 |
| S8000047^T^ | C1=C(C3=C(C(=N1)N[C@@H](C(F)(F)F)C2CC2)C4=C([NH]3)C=C(C=C4)C5=CN=C(N=C5)N6CCOCC6)C(=O)N | 0.903 | 0.1783 |
| S8000048^P^ | C1=C(C3=C(C(=N1)N[C@@H](C(F)(F)F)C2CC2)C4=C([NH]3)C=C(C=C4)C5=NN=C(C=C5)N6CC[S](CC6)(=O)=O)C(=O)N | 0.301 | 0.1499 |
| S8000049^T^ | C1=C(C3=C(C(=N1)N[C@@H](C(F)(F)F)C2CC2)C4=C([NH]3)C=C(C=C4)C5=CN=C(N=C5)N)C(=O)N | -0.097 | 0.2092 |

#T: Represents compounds present in training set.

#P: Represents compounds classified as prediction set.

\# Exp-log IC_50 :_ Experimental Log IC_50_ values obtained from literature.

\# pIC_50_ **_:_** Predicted Log IC_50_ values obtained by Multi-Linear Regression (MLR) based calculation.

**Table S2**: Represents significant statistical parameters associated with validated QSAR model consisting of 3 descriptors.

| **Compound ID** | **Status** | **Pred.Mod.Eq.Res** | **Pred.LOO** | **Pred.LOO.Res** | **HAT i/i(h*= 0.3429)** |
| --- | --- | --- | --- | --- | --- |
| S8000001 | Training | -0.0358 | 2.271 | -0.051 | 0.2979 |
| S8000002 | Training | -0.2388 | 2.0722 | -0.2898 | 0.1758 |
| S8000003 | Prediction | 0.009 | - | - | 0.0818 |
| S8000004 | Prediction | -0.0352 | - | - | 0.0584 |
| S8000005 | Training | 0.1352 | 2.1577 | 0.1577 | 0.1426 |
| S8000006 | Training | 0.0139 | 1.8839 | 0.0149 | 0.0699 |
| S8000007 | Training | 0.0497 | 1.7684 | 0.0524 | 0.0523 |
| S8000008 | Training | -0.5563 | 1.7349 | -0.5871 | 0.0523 |
| S8000009 | Training | 0.3307 | 2.2982 | 0.3792 | 0.128 |
| S8000010 | Training | 0.0347 | 1.7461 | 0.0381 | 0.0874 |
| S8000011 | Training | 0.3487 | 1.607 | 0.377 | 0.0749 |
| S8000012 | Training | 0.0931 | 1.3808 | 0.1018 | 0.0859 |
| S8000013 | Training | 0.4079 | 1.6862 | 0.4312 | 0.054 |
| S8000014 | Training | -0.4883 | 1.7981 | -0.5639 | 0.134 |
| S8000015 | Prediction | -0.4785 | - | - | 0.0565 |
| S8000017 | Training | -0.0333 | 1.7342 | -0.0368 | 0.0942 |
| S8000018 | Training | 0.3981 | 1.8043 | 0.4423 | 0.1001 |
| S8000019 | Training | -0.344 | 1.6574 | -0.3836 | 0.103 |
| S8000020 | Prediction | 0.2096 | - | - | 0.2487 |
| S8000021 | Training | -0.2129 | 1.9715 | -0.2835 | 0.249 |
| S8000022 | Prediction | 0.4781 | - | - | 0.0353 |
| S8000023 | Prediction | 0.0606 | - | - | 0.0353 |
| S8000025 | Training | -0.1903 | 1.1651 | -0.1969 | 0.0333 |
| S8000026 | Training | 0.3595 | 1.0704 | 0.3714 | 0.0321 |
| S8000027 | Training | -0.5499 | 1.0363 | -0.5767 | 0.0464 |
| S8000028 | Training | 0.1589 | 1.259 | 0.18 | 0.1169 |
| S8000029 | Training | 0.1589 | 1.259 | 0.18 | 0.1169 |
| S8000030 | Prediction | 0.382 | - | - | 0.0897 |
| S8000031 | Training | 0.208 | 1.2377 | 0.2377 | 0.1249 |
| S8000032 | Prediction | 0.6672 | - | - | 0.0863 |
| S8000033 | Prediction | -0.3738 | - | - | 0.0863 |
| S8000034 | Training | 0.247 | 0.5703 | 0.2693 | 0.0827 |
| S8000035 | Training | 0.081 | 0.566 | 0.089 | 0.0898 |
| S8000036 | Training | 0.0659 | 0.3837 | 0.0827 | 0.2024 |
| S8000037 | Training | 0.6021 | 0.2805 | 0.6785 | 0.1126 |
| S8000038 | Training | -0.7499 | 0.109 | -0.845 | 0.1126 |
| S8000039 | Prediction | 0.0824 | - | - | 0.143 |
| S8000040 | Training | -0.4285 | 0.6052 | -0.4738 | 0.0957 |
| S8000041 | Prediction | -0.0998 | - | - | 0.1788 |
| S8000042 | Training | 0.2841 | 0.6555 | 0.3545 | 0.1987 |
| S8000043 | Training | -0.3218 | 0.238 | -0.364 | 0.116 |
| S8000044 | Training | 0.3178 | 0.3601 | 0.3601 | 0.1175 |
| S8000045 | Training | 0.1562 | 0.1816 | 0.1816 | 0.14 |
| S8000046 | Training | 0.1172 | 0.4284 | 0.1274 | 0.0799 |
| S8000047 | Training | -0.7247 | 0.0817 | -0.8213 | 0.1176 |
| S8000048 | Prediction | -0.1511 | - | - | 0.1469 |
| S8000049 | Training | 0.3062 | 0.2686 | 0.3656 | 0.1626 |

#Pred.Mod.Eq. Res: Residual values calculated from the predicted IC50 values obtained using model equation.

#Pred.LOO: Values predicted by Leave One Out method (internal validation).

#Pred.LOO.Res: Residual values calculated from the predicted IC50 values obtained by performing Leave One Out method.

#HAT: Leverage value taken from HAT matrix diagonal.

**Table S3**: Details of the molecular descriptor values present in validated QSAR model.

| **ID** | **Average broto-moreau autocorrelation - lag 1/weighted by first ionization potential (D1)** | **Largest absolute Eigen value of burden modified matrix-n7/weighted by relative polarizability (D2)** | **Self returning walk count of order 9 (D3)** |
| --- | --- | --- | --- |
| S8000001 | 152.472 | 2.315 | 7.081 |
| S8000002 | 151.775 | 2.425 | 7.081 |
| S8000003 | 151.189 | 2.575 | 7.081 |
| S8000004 | 150.688 | 2.682 | 7.081 |
| S8000005 | 151.189 | 2.458 | 7.081 |
| S8000006 | 150.688 | 2.591 | 7.081 |
| S8000007 | 149.878 | 2.69 | 7.081 |
| S8000008 | 149.878 | 2.69 | 7.081 |
| S8000009 | 149.969 | 2.491 | 7.081 |
| S8000010 | 149.577 | 2.617 | 7.445 |
| S8000011 | 149.236 | 2.806 | 7.081 |
| S8000012 | 148.936 | 2.876 | 7.195 |
| S8000013 | 150.266 | 2.706 | 7.081 |
| S8000014 | 149.778 | 2.549 | 7.455 |
| S8000015 | 149.399 | 2.707 | 7.081 |
| S8000017 | 148.299 | 2.803 | 7.081 |
| S8000018 | 148.145 | 2.804 | 7.081 |
| S8000019 | 148.145 | 2.825 | 7.096 |
| S8000020 | 145.997 | 2.825 | 7.081 |
| S8000021 | 145.997 | 2.83 | 7.081 |
| S8000022 | 149.419 | 2.825 | 7.545 |
| S8000023 | 149.419 | 2.826 | 7.545 |
| S8000025 | 149.842 | 2.825 | 7.455 |
| S8000026 | 149.936 | 2.839 | 7.545 |
| S8000027 | 149.186 | 2.883 | 7.554 |
| S8000028 | 150.833 | 2.839 | 7.081 |
| S8000029 | 150.833 | 2.839 | 7.081 |
| S8000030 | 149.038 | 2.839 | 7.081 |
| S8000031 | 151.018 | 2.839 | 7.081 |
| S8000032 | 149.7 | 2.858 | 8.091 |
| S8000033 | 149.7 | 2.858 | 8.091 |
| S8000034 | 149.419 | 2.922 | 8.097 |
| S8000035 | 149.165 | 2.933 | 8.102 |
| S8000036 | 148.652 | 2.943 | 8.458 |
| S8000037 | 152.093 | 2.884 | 8.107 |
| S8000038 | 152.093 | 2.884 | 8.107 |
| S8000039 | 152.59 | 2.867 | 7.545 |
| S8000040 | 151.685 | 2.889 | 7.545 |
| S8000041 | 149.019 | 2.925 | 8.423 |
| S8000042 | 147.848 | 2.953 | 8.296 |
| S8000043 | 149.533 | 2.966 | 8.296 |
| S8000044 | 149.533 | 2.951 | 8.296 |
| S8000045 | 152.699 | 2.878 | 8.057 |
| S8000046 | 151.716 | 2.889 | 7.87 |
| S8000047 | 151.247 | 3.015 | 7.87 |
| S8000048 | 150.027 | 3.105 | 7.87 |
| S8000049 | 153.056 | 2.886 | 7.87 |

| 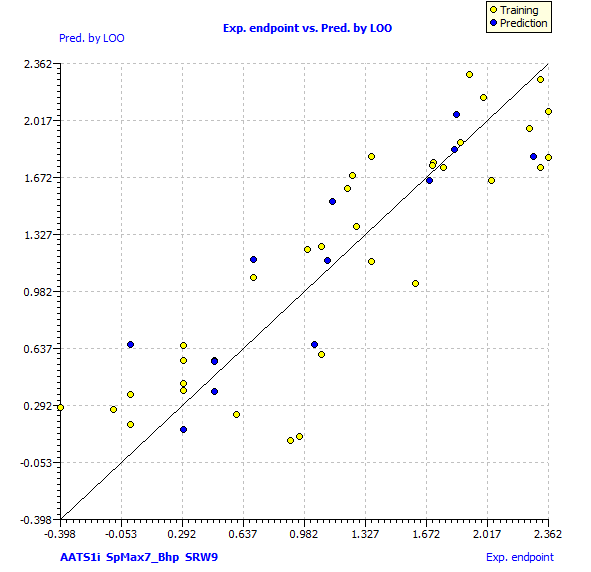 | 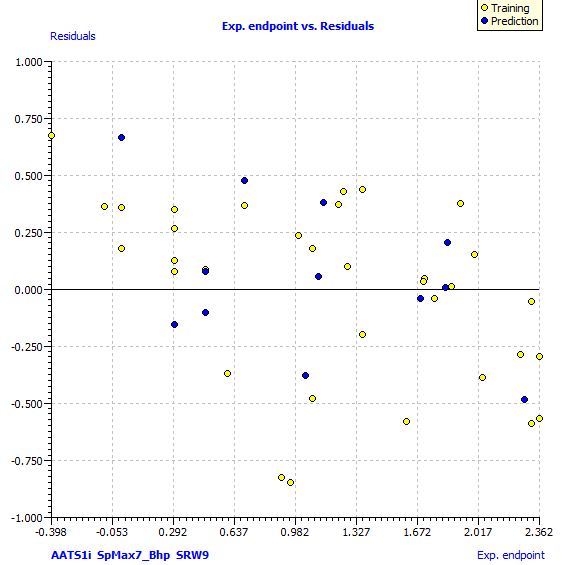 |
| --- | --- |

**Figure S2**: Residual plot showing the relationship between experimental Log IC50 and residual values. Residual values are calculated by taking the difference of experimental and predicted Log IC_50_.

**Figure S1**: Scatter plot depicting the relationship between experimental and predicted Log IC_50_. The predicted Log IC_50_ is calculated by performing Leave One Out (LOO) internal validation method, were single observation is considered for validation and rest of the data points in the dataset are considered as training data for model building.

| 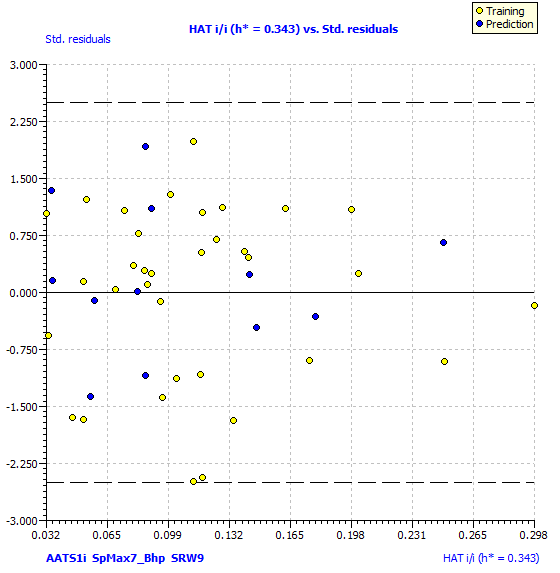 | 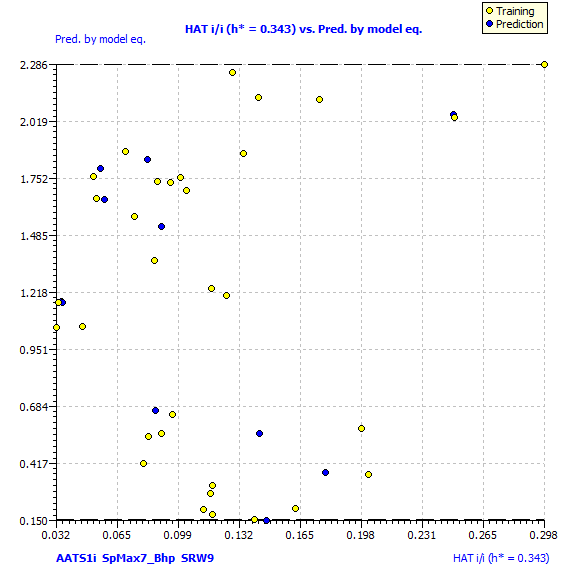 |
| --- | --- |

**Figure S4**: Insubria graph showing the relationship between HAT values and predicted Log IC_50_ values.

**Figure S3**: Depicts William’s plot. The plot of standard residuals and HAT values provide significant information on Applicability Domain (AD) of QSAR model. The presence of data points within AD shows the structural similarity of chemical compounds used for model building and the reliability of model in predicting activities of new compounds.

**Table S4**: Provides significant information regarding compounds used for external validation of QSAR model and their descriptor values.

| **Sl.no** | **ID** | **Structure (SMILES)** | **D1** | **D2** | **D3** | **pIC_50_** | **logIC_50_** |
| --- | --- | --- | --- | --- | --- | --- | --- |
| 1 | S2000001 | N#CC[C@H](C1CCCC1)N1C=C(C=N1)C1=NC=NC2=C1C=CN2 | 148.87 | 2.54 | 7.60 | 1.91 | 0.44 |
| 2 | S2000002 | CC1=C(NC2=CC=CC(=C2)S(=O)(=O)NC(C)(C)C)N=C(NC2=CC=C(OCCN3CCCC3)C=C2)N=C1 | 148.09 | 3.11 | 6.25 | 1.55 | 0.47 |
| 3 | S2000003 | O=C(NCC#N)C1=CC=C(C=C1)C1=NC(NC2=CC=C(C=C2)N2CCOCC2)=NC=C1 | 148.04 | 2.82 | 0 | 6.67 | 3.30 |
| 4 | S2000004 | C[C@H](NC(=O)C(=C\C1=NC(Br)=CC=C1)\C#N)C1=CC=CC=C1 | 143.82 | 2.65 | 0 | 7.78 | 3.36 |
| 5 | S2000005 | CN1CCN(CC1)C1=CC=C(NC2=NC=C(C)C(NC3=CC(=CC=C3)S(=O)(=O)NC(C)(C)C)=N2)C=C1 | 149.09 | 2.97 | 0 | 6.13 | 0.77 |

\# pIC_50_ – Predicted IC50 values calculated by using MLR equation.

\# log IC_50_ – Logarithm of IC50 values obtained from literature.

**Table S5**: Provides vital information regarding dock score, binding affinity and ligand efficiency of compounds used for external validation with JAK2 protein.

| **Compound** | **Dock score** | **Binding energy**  **(kJ/mol)** | **Ligand efficiency** | **Binding affinity range** |
| --- | --- | --- | --- | --- |
| S2000001 | -28.51 | -31 | 0.33 | μM-nM |
| S2000002 | -30.49 | -12 | 0.08 | mM |
| S2000003 | -25.94 | -10 | 0.08 | Mm-μM |
| S2000004 | -21.80 | -26 | 0.28 | μM-nM |
| S2000005 | -27.94 | -21 | 0.14 | μM-nM |

| **S2000001 Dock score -28.51 Energy -31 kJ/mol**  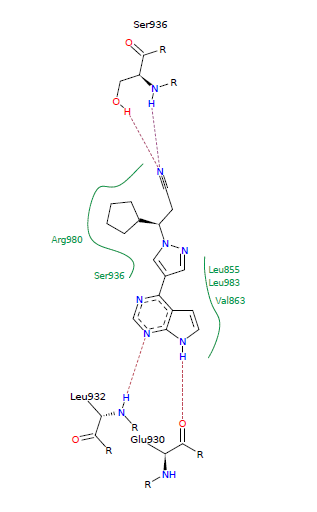  **a.** | **S2000002 Dock score -30.49 Energy -12 kJ/mol**  **b.** 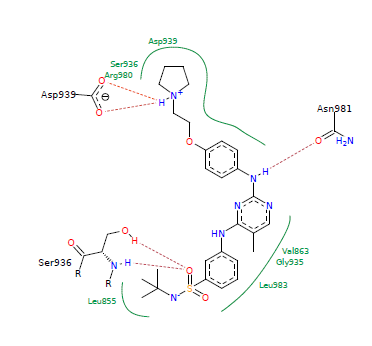 |
| --- | --- |

| **S2000003 Dock score -25.94 Energy -10kJ/mol**  **c**. 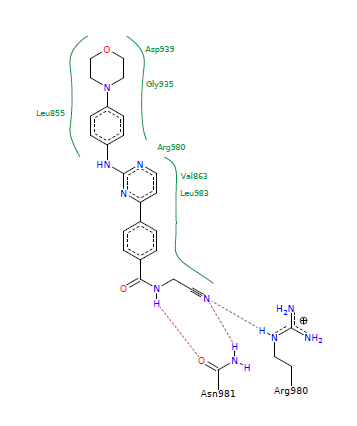 | **S2000004 Dock score -21.80 Energy -26kJ/mol**  **d.** 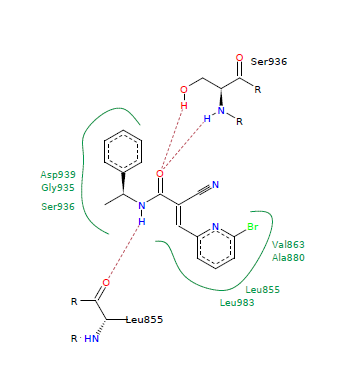 |
| --- | --- |

**S2000005	Dock score -27.94	Energy -21kJ/mol**

**e**.
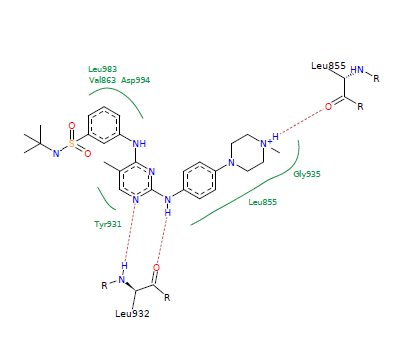

**Figure S5**: Depicts binding interactions of compounds used for external validation of QSAR model with JAK2 protein. Dock score is calculated by considering contributions of lipophilic contact energy, energy derived from hydrogen bond interactions, ionization energies and number of rotatable bonds. Binding energy is calculated by considering desolvation parameters. Negative values of dock score and binding energy provides insights into the comfortability of ligand to interact with the protein and stability of protein-ligand complex respectively.

**Table S6**: Provides significant information regarding dock score, binding affinity and ligand efficiency of all compounds present in the dataset.

| **Compound ID** | **Dock score** | **Binding energy (kJ/mol)** | **Ligand efficiency** | **Binding affinity range**  **(mM-nM)** |
| --- | --- | --- | --- | --- |
| S8000001 | -31.66 | -31 | 0.39 | mM - μM |
| S8000002 | -27.74 | -16 | 0.19 | mM-μM |
| S8000003 | -27.84 | -15 | 0.17 | mM-μM |
| S8000004 | -25.91 | -12 | 0.13 | mM-μM |
| S8000005 | -28.84 | -14 | 0.16 | mM-μM |
| S8000006 | -28.84 | -11 | 0.12 | mM-μM |
| S8000007 | -26.58 | -10 | 0.1 | mM-μM |
| S8000008 | -27.15 | -27 | 0.27 | μM |
| S8000009 | -29.95 | -13 | 0.14 | mM-μM |
| S8000010 | -30.93 | -11 | 0.11 | mM-μM |
| S8000011 | -28.72 | -11 | 0.11 | mM-μM |
| S8000012 | -30.18 | -15 | 0.14 | mM-μM |
| S8000013 | -33.01 | -13 | 0.13 | mM-μM |
| S8000014 | -26.54 | -14 | 0.16 | mM-μM |
| S8000015 | -27.59 | -14 | 0.15 | mM-μM |
| S8000017 | -29.12 | -13 | 0.13 | mM-μM |
| S8000018 | -27.47 | -12 | 0.12 | mM-μM |
| S8000019 | -29.72 | -15 | 0.15 | mM-μM |
| S8000020 | -28.28 | -3 | 0.03 | mM-μM |
| S8000021 | -28.45 | -15 | 0.12 | mM-μM |
| S8000022 | -29.26 | -8 | 0.07 | mM-μM |
| S8000023 | -30.11 | -8 | 0.07 | mM-μM |
| S8000025 | -31.2 | 12 | 0 | mM-μM |
| S8000026 | -26.9 | -19 | 0.16 | μM |
| S8000027 | -24.31 | -18 | 0.13 | mM-μM |
| S8000028 | -28.81 | -2 | 0.02 | mM-μM |
| S8000029 | -31.26 | -23 | 0.18 | mM-μM |
| S8000030 | -28.74 | 20 | 0 | mM-μM |
| S8000031 | -29.34 | 15 | 0 | mM-μM |
| S8000032 | -26.33 | -37 | 0.31 | nM |
| S8000033 | -26.44 | -32 | 0.27 | μM |
| S8000034 | -25.67 | 11 | 0 | mM-μM |
| S8000035 | -25.51 | -34 | 0.27 | μM |
| S8000036 | -25.31 | -34 | 0.27 | μM |
| S8000037 | -25.27 | -25 | 0.19 | μM |
| S8000038 | -25.44 | -26 | 0.2 | μM |
| S8000039 | -26.56 | -27 | 0.21 | μM |
| S8000040 | -25.01 | -9 | 0.07 | mM-μM |
| S8000041 | -27.17 | -34 | 0.28 | nM |
| S8000042 | -26.9 | -35 | 0.28 | nM |
| S8000043 | -27.08 | -26 | 0.2 | μM |
| S8000044 | -26.84 | -9 | 0.07 | mM-μM |
| S8000045 | -29.99 | -23 | 0.18 | μM |
| S8000046 | -25.46 | -30 | 0.22 | μM |
| S8000047 | -26.85 | -15 | 0.1 | mM-μM |
| S8000048 | -26.21 | 12 | 0 | mM-μM |
| S8000049 | -26.09 | -34 | 0.25 | μM |

\# Ligand efficiency depicts the measurement of binding energy per atom to it’s binding partner.

\# Binding affinity in nM range is highly acceptable for protein-ligand complexes.

| 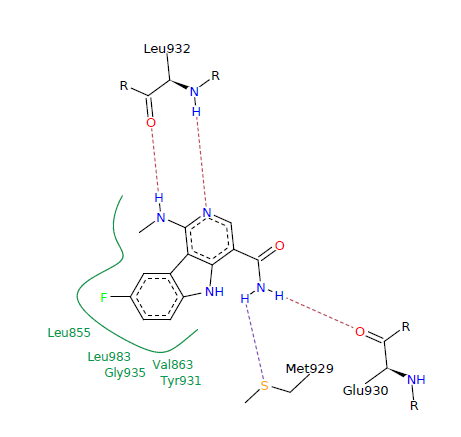**S8000001 Dock score: -31.66 Energy: -31 kJ/mol** | **S8000002 Dock score: -27.74 Energy: -16 kJ/mol**  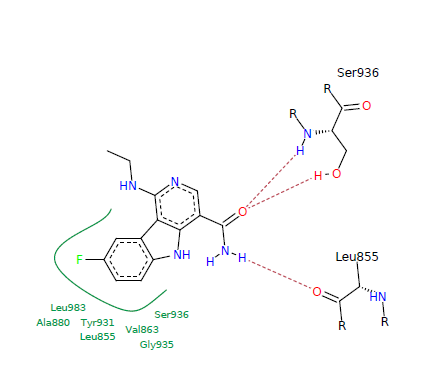 |
| --- | --- |
| **S8000003 Dock score: -27.84 Energy: -15 kJ/mol**  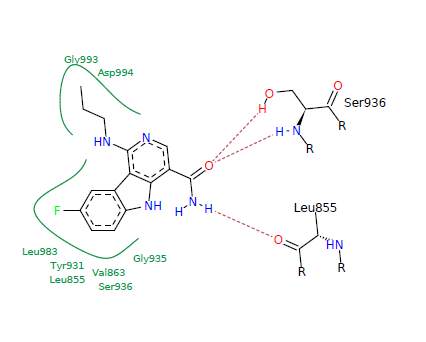 | **S8000004 Dock score: -25.91 Energy: -12 kJ/mol**  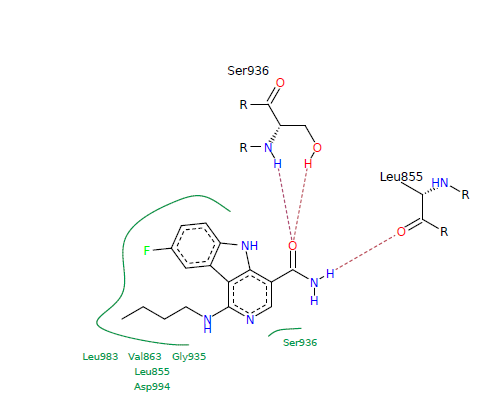 |
| 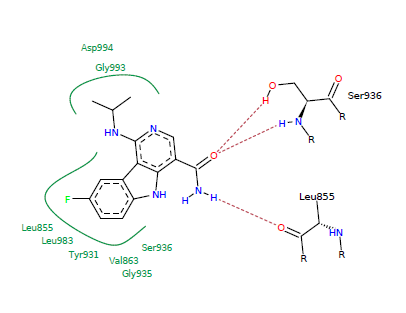**S8000005 Dock score: -28.84 Energy: -14** **kJ/mol** | 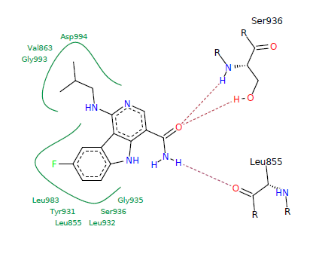**S8000006 Dock score: -28.84 Energy: -11** **kJ/mol** |
| 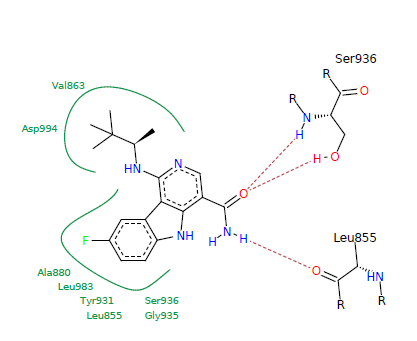**S8000007 Dock score: -26.58 Energy: -10 kJ/mol** | 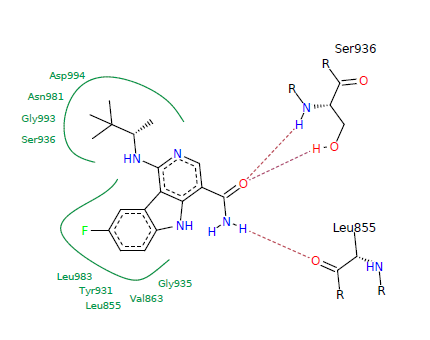**S8000008 Dock score: -27.15 Energy: -27** **kJ/mol** |
| 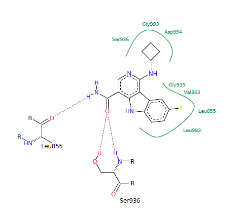**S8000009 Dock score: -29.95 Energy: -13 kJ/mol** | 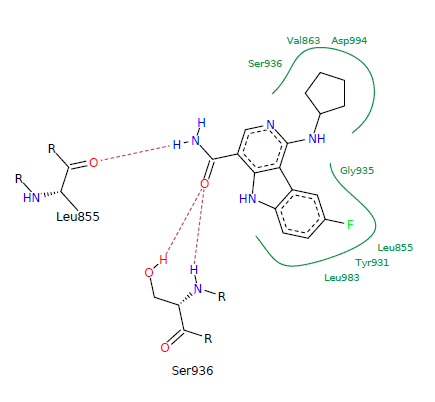**S8000010 Dock score: -30.93 Energy: -11** **kJ/mol** |
| **S8000011 Dock score: -28.72 Energy: -11** **kJ/mol**  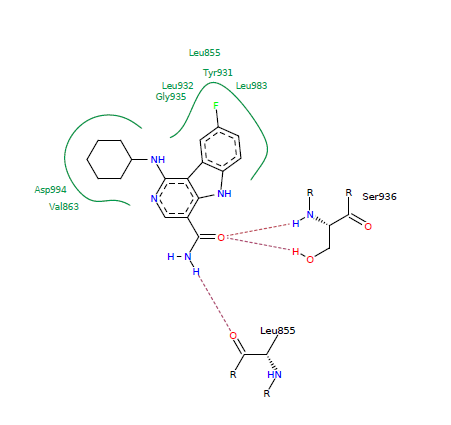 | 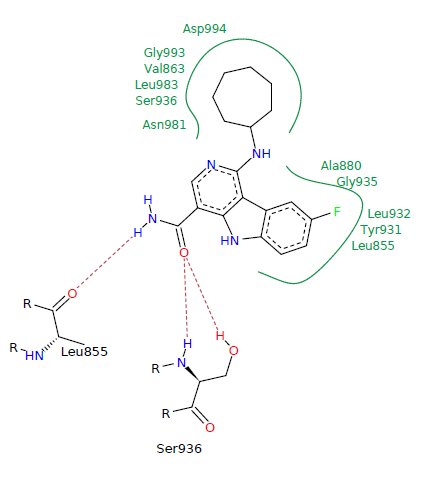**S8000012 Dock score: -30.18 Energy: -15** **kJ/mol** |
| 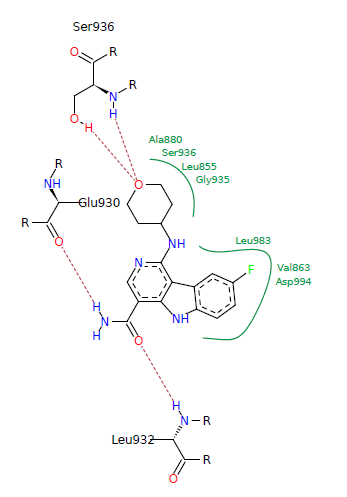**S8000013 Dock score: -33.01 Energy: -13** **kJ/mol** | 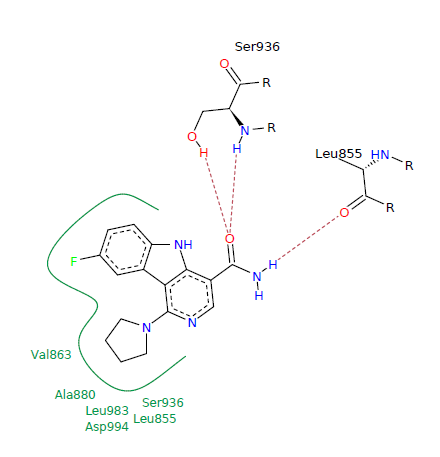**S8000014 Dock score: -26.54 Energy: -14** **kJ/mol** |
| **S8000015 Dock score: -27.59 Energy: -14** **kJ/mol**  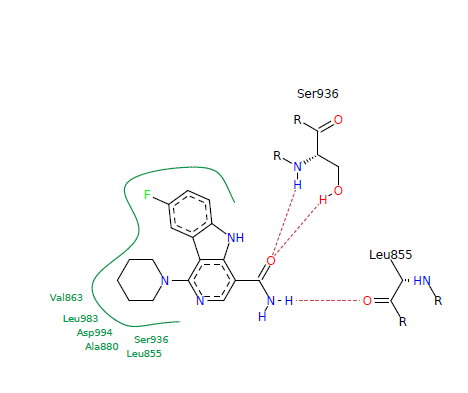 | **S8000017 Dock score: -29.12 Energy: -13** **kJ/mol**  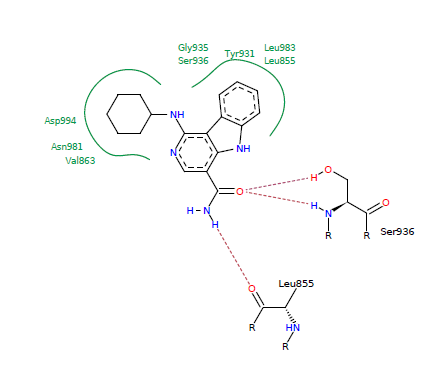 |
| 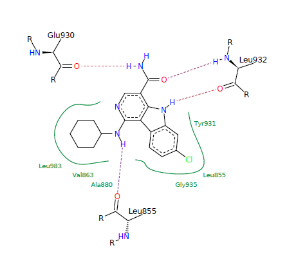**S8000018 Dock score: -27.47 Energy: -12** **kJ/mol** | 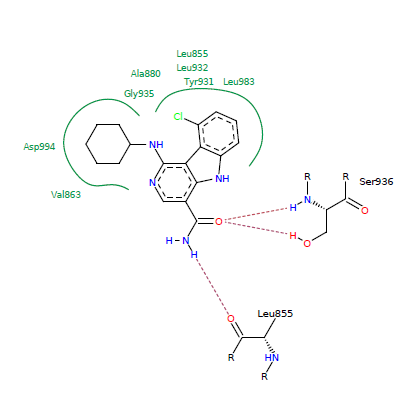**S8000019 Dock score: -29.72 Energy: -15** **kJ/mol** |
| 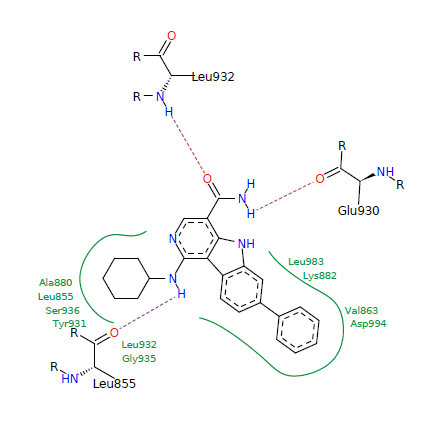**S8000020 Dock score: -28.28 Energy: -3** **kJ/mol** | 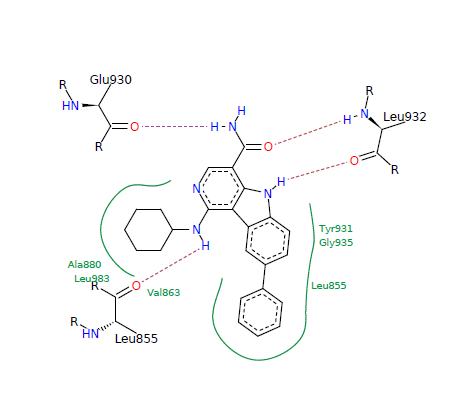**S8000021 Dock score: -28.45 Energy: -15** **kJ/mol** |
| 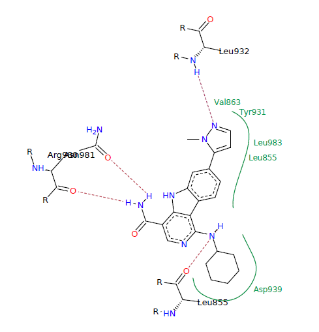**S8000022 Dock score: -29.26 Energy: -8** **kJ/mol** | 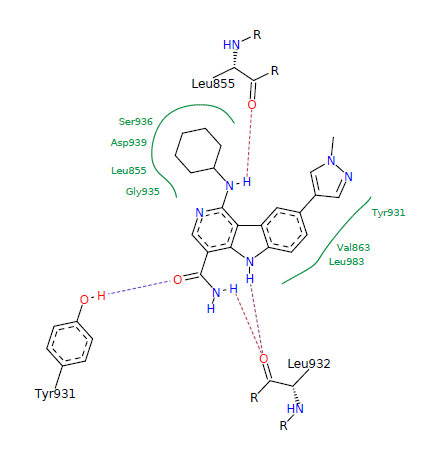**S8000023 Dock score: -30.11 Energy: -8** **kJ/mol** |

| 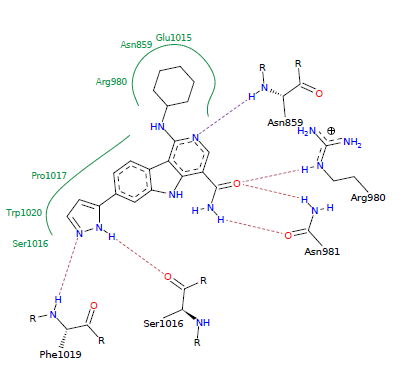**S8000025 Dock score: -31.2 Energy: 12** **kJ/mol** | 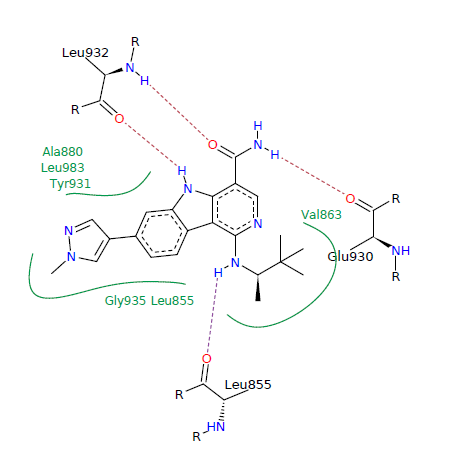**S8000026 Dock score: -26.9 Energy: -19** **kJ/mol** | |
| --- | --- | --- |
| 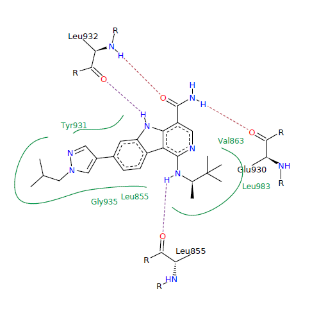**S8000027 Dock score: -24.31 Energy: -18** **kJ/mol** | 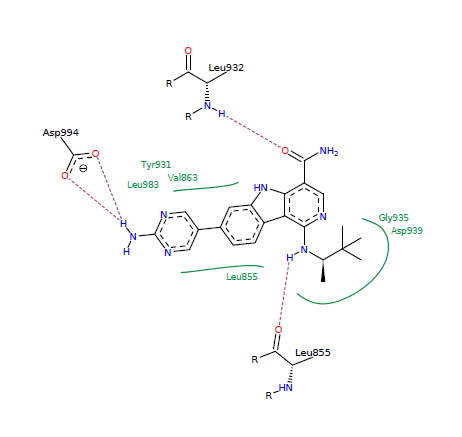**S8000028 Dock score: -28.81 Energy: -2** **kJ/mol** | |
| **S8000029 Dock score: -31.26 Energy: -23** **kJ/mol**  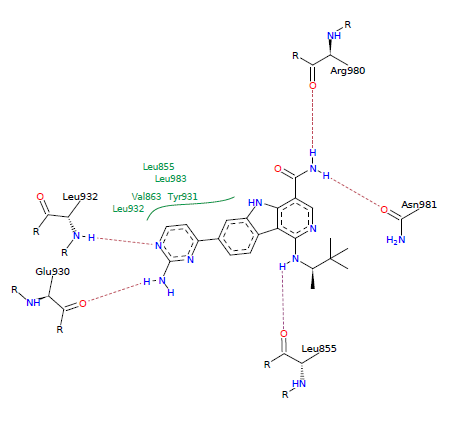 | | **S8000030 Dock score: -28.74 Energy: 20** **kJ/mol**  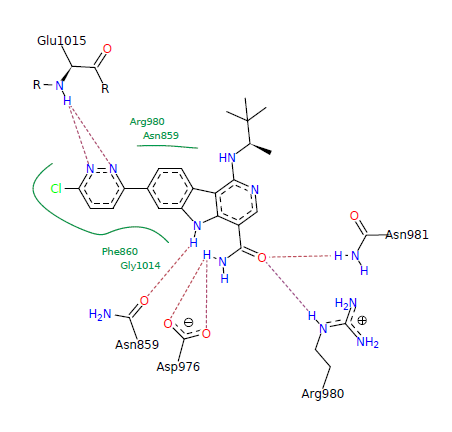 |
| 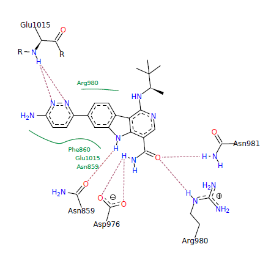**S8000031 Dock score: -29.34 Energy: 15** **kJ/mol** | | 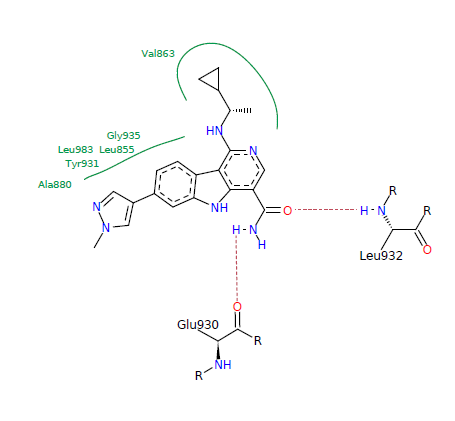**S8000032 Dock score: -26.33 Energy: -37** **kJ/mol** |
| **S8000033 Dock score: -26.44 Energy: -32** **kJ/mol**  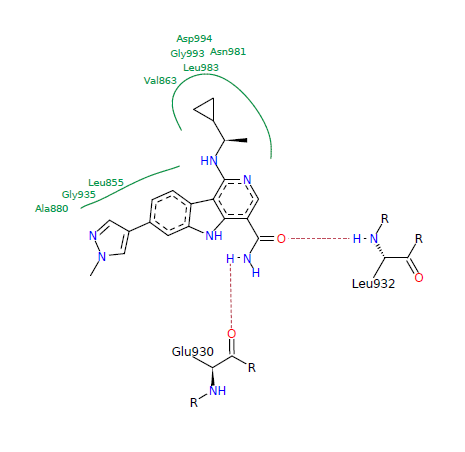 | | **S8000034 Dock score: -25.67 Energy: 11** **kJ/mol**  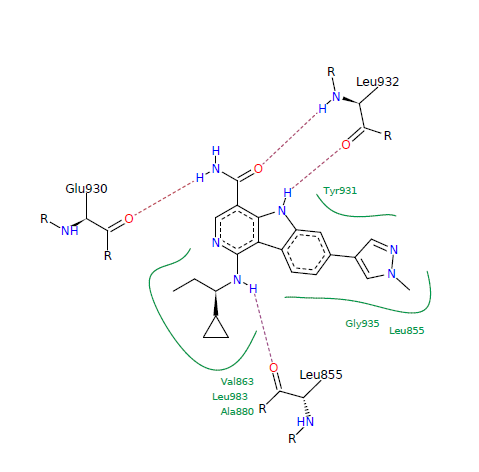 |
| 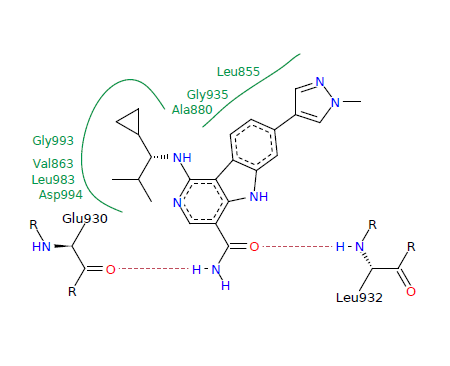**S8000035 Dock score: -25.51 Energy: -34** **kJ/mol** | | 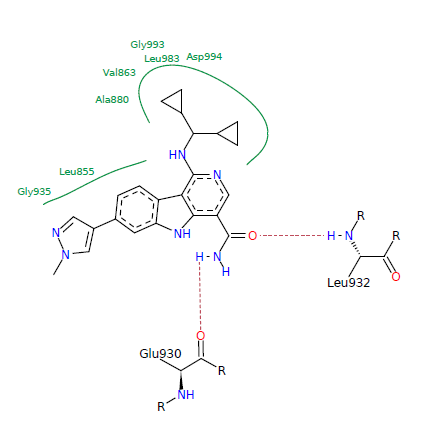**S8000036 Dock score: -25.31 Energy: -34** **kJ/mol** |
| 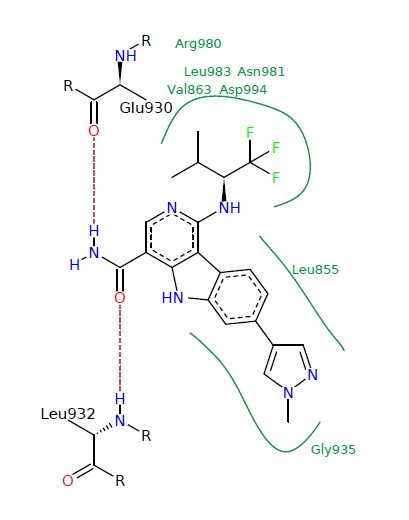**S8000039 Dock score: -26.56 Energy: -27** **kJ/mol** | | **S8000040 Dock score: -25.01 Energy: -9** **kJ/mol**  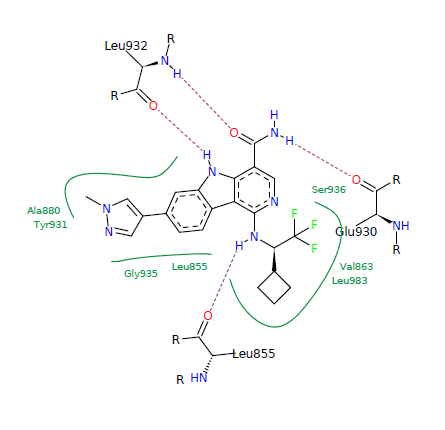 |
| 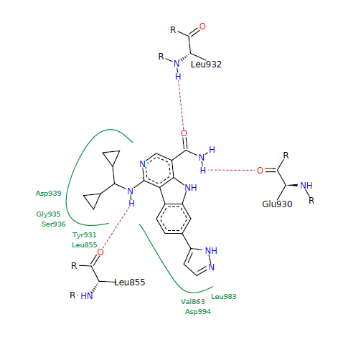**S8000041 Dock score: -27.17 Energy: -34** **kJ/mol** | | **S8000042 Dock score: -26.9 Energy: -35** **kJ/mol**  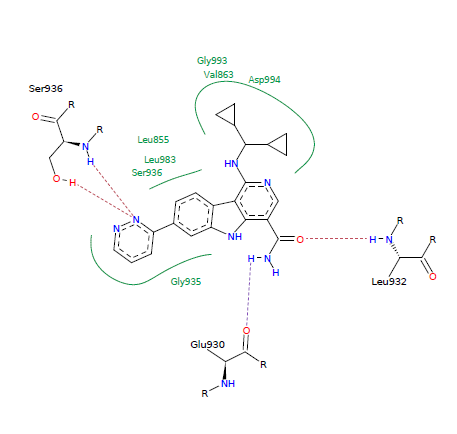 |
| **S8000043 Dock score: -27.08 Energy: -26** **kJ/mol**  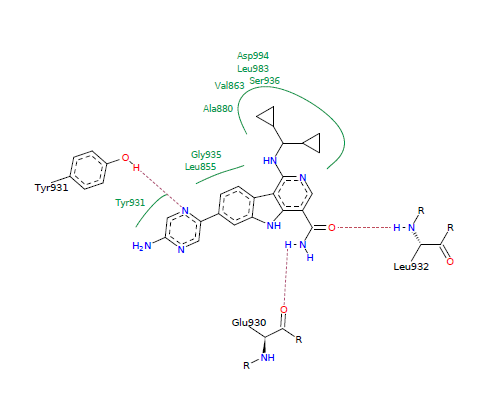 | | **S8000044 Dock score: -26.84 Energy: -9** **kJ/mol**  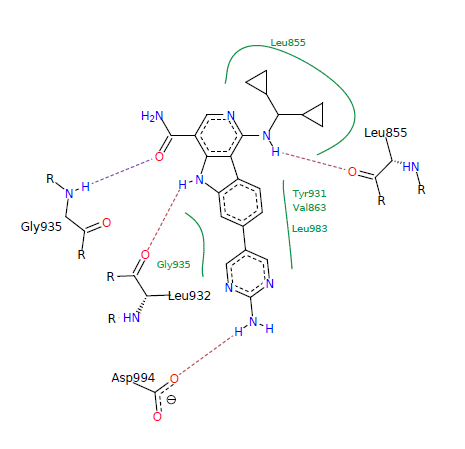 |
| 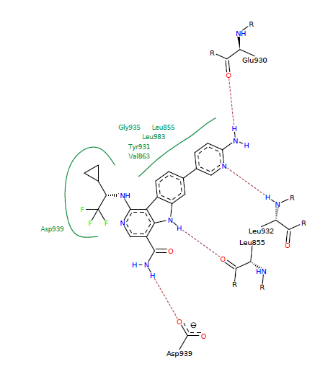**S8000046 Dock score: -25.46 Energy: -30** **kJ/mol** | | 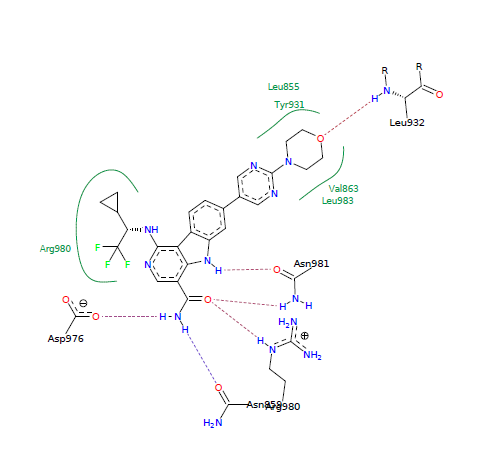**S8000047 Dock score: -26.85 Energy: -15** **kJ/mol** |
| **S8000048 Dock score: -26.21 Energy: 12** **kJ/mol**  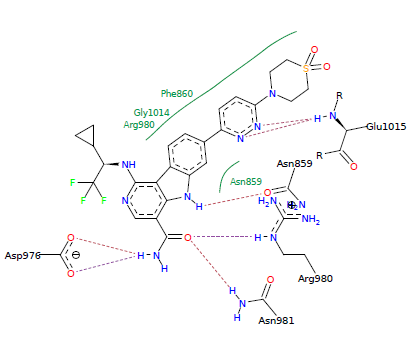 | | **S8000049 Dock score: -26.09 Energy: -34** **kJ/mol**  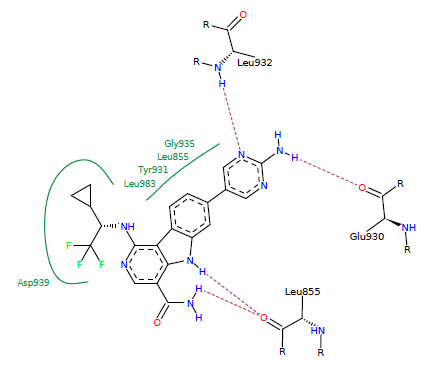 |

**Figure S6 :** Represents binding interactions of all compounds present in dataset used for QSAR modelling with JAK2 protein.

**Table S7**: Provides significant information regarding factors effecting Absorption, Distribution, Metabolism, Elimination (ADME) (bioavailability) parameters for all the compounds used for QSAR model building.

| **Compound ID** | **logS** | **logP** | **2C9 pKi** | **hERG pIC_50_** | **HIA category** | **2D6 affinity category** | **PPB90 category** |
| --- | --- | --- | --- | --- | --- | --- | --- |
| S8000001 | 1.912 | 1.485 | 4.808 | 4.194 | + | low | low |
| S8000002 | 1.509 | 1.919 | 5.012 | 4.331 | + | medium | low |
| S8000003 | 1.376 | 2.312 | 5.113 | 4.515 | + | medium | low |
| S8000004 | 1.251 | 2.69 | 5.225 | 4.667 | + | medium | high |
| S8000005 | 1.626 | 2.374 | 5.119 | 4.425 | + | high | low |
| S8000006 | 1.27 | 2.637 | 5.192 | 4.518 | + | high | high |
| S8000007 | 1.236 | 3.58 | 5.409 | 4.689 | + | medium | high |
| S8000008 | 1.236 | 3.58 | 5.409 | 4.689 | + | medium | high |
| S8000009 | 1.36 | 2.47 | 5.103 | 4.663 | + | high | low |
| S8000010 | 1.185 | 2.802 | 4.932 | 4.784 | + | high | high |
| S8000011 | 1.017 | 3.33 | 5.051 | 4.927 | + | high | high |
| S8000012 | 0.8771 | 3.769 | 5.048 | 5.06 | + | very high | high |
| S8000013 | 1.241 | 2.01 | 5.043 | 4.623 | + | high | low |
| S8000014 | 0.8003 | 2.235 | 5.294 | 4.757 | + | medium | low |
| S8000015 | 0.634 | 2.808 | 5.424 | 4.911 | + | medium | low |
| S8000016 | 0.6049 | 4.114 | 5.082 | 5.215 | + | very high | high |
| S8000017 | 0.9478 | 3.274 | 4.93 | 4.695 | + | high | low |
| S8000018 | 0.9276 | 3.851 | 5.131 | 4.725 | + | high | high |
| S8000019 | 0.9192 | 3.902 | 5.055 | 4.778 | + | high | high |
| S8000020 | 0.4055 | 4.682 | 5.328 | 5.001 | + | high | high |
| S8000021 | 0.4055 | 4.682 | 5.328 | 4.998 | + | high | high |
| S8000022 | 1.342 | 3.929 | 5.256 | 4.931 | + | high | high |
| S8000023 | 1.324 | 3.929 | 5.213 | 4.926 | + | high | high |
| S8000024 | 1.246 | 3.091 | 5.2 | 4.334 | + | high | high |
| S8000025 | 1.246 | 3.091 | 5.2 | 4.334 | + | high | high |
| S8000026 | 1.458 | 4.128 | 5.714 | 4.682 | + | medium | high |
| S8000027 | 1.112 | 4.955 | 5.898 | 4.963 | + | high | high |
| S8000028 | 1.249 | 3.504 | 5.894 | 4.382 | + | medium | high |
| S8000029 | 1.284 | 3.504 | 5.925 | 4.319 | + | medium | high |
| S8000030 | 1.091 | 4.679 | 5.951 | 4.74 | + | medium | high |
| S8000031 | 1.298 | 3.422 | 5.9 | 4.383 | + | medium | high |
| S8000032 | 1.29 | 3.46 | 5.631 | 4.676 | + | high | high |
| S8000033 | 1.602 | 3.46 | 5.631 | 4.676 | + | high | high |
| S8000034 | 1.479 | 3.805 | 5.628 | 4.804 | + | high | high |
| S8000035 | 1.366 | 4.068 | 5.658 | 4.838 | + | very high | high |
| S8000036 | 1.381 | 3.88 | 5.276 | 4.953 | + | very high | high |
| S8000037 | 1.581 | 3.603 | 5.879 | 4.991 | + | medium | high |
| S8000038 | 1.581 | 3.603 | 5.879 | 4.991 | + | medium | high |
| S8000039 | 1.581 | 3.76 | 5.879 | 4.903 | + | medium | high |
| S8000040 | 1.416 | 3.914 | 5.843 | 5.122 | + | medium | high |
| S8000041 | 1.416 | 3.018 | 5.193 | 4.359 | + | high | high |
| S8000042 | 1.337 | 3.887 | 5.306 | 4.915 | + | very high | high |
| S8000043 | 1.153 | 3.182 | 5.426 | 4.652 | - | very high | high |
| S8000044 | 1.104 | 3.288 | 5.427 | 4.651 | - | very high | high |
| S8000045 | 1.494 | 2.867 | 5.706 | 4.391 | + | medium | high |
| S8000046 | 1.507 | 2.754 | 6.046 | 4.536 | + | medium | high |
| S8000047 | 0.2942 | 3.73 | 6.023 | 5.477 | + | medium | high |
| S8000048 | 1.055 | 2.711 | 6.054 | 5.456 | + | medium | high |
| S8000049 | 1.295 | 3.227 | 5.979 | 4.701 | - | medium | high |

**Table S8:** Details regarding structure and physicochemical properties of derivative compounds identified for compound S8000041.

| **Structure (SMILES)** | **MW** | **HBD** | **HBA** | **TPSA** | **Rotatable Bonds** |
| --- | --- | --- | --- | --- | --- |
| O=C(NC1CC[NH+](C(C)(C)C)CC1)NCC2=CC=C(C=C2)C3=N[NH]C=C3 (**I1**) | 355.5 | 3 | 6 | 73.05 | 7 |
| FC2=C([N]1C=NC=C1)C=CC(=C2)CNC(=O)C(=O)N[C@H]4C[NH+](C3CC3)CC4 (**I2**) | 371.4 | 2 | 7 | 79.26 | 8 |
| O=C(N2C[C@H]([NH+](C1CC1)CC2)C)C(=O)NCC3=C4C(=CC=C3)N=CC=C4 **(I3**) | 352.4 | 1 | 6 | 65.54 | 6 |
| O=C(NCC2=CC1=C([C](C=C[NH]1)=O)C=C2)NC[C@H]3C[NH+](CC(C)C)CC3 **(I4**) | 356.5 | 3 | 6 | 77.23 | 8 |
| O=C(NNC1=CC2=C(C=C1)[C](=O)[NH]C=C2)NC4=C(NC3CC3)C=CC(=C4)C(=O)N (**I5**) | 392.4 | 6 | 9 | 141.1 | 8 |
| FC2=C([N]1C=NC=C1)C=CC(=C2)CNC(=O)C(=O)N[C@H]4[C@@H](C[NH+](C3CC3)C4)C **(I6**) | 385.4 | 2 | 7 | 79.26 | 8 |
| O=C(N2CC[NH+](C1CC1)CC2)C(=O)NCC3=C4C(=CC=C3)N=CC=C4 (**I7**) | 338.4 | 1 | 6 | 65.54 | 6 |
| O=C(NC2CC[NH+](CC1CC1)CC2)NCC3=CC=C(C=C3)C4=N[NH]C=C4 (**I8**) | 353.5 | 3 | 6 | 73.05 | 8 |
| O=C(NCC1=CC=C(C=C1)C2=N[NH]C=C2)NCC4CC[NH+](CC3CC3)CC4 (**I9**) | 367.5 | 3 | 6 | 73.05 | 9 |
| BrC1=C2C(=CC=C1)[NH]C=C2C[C@H]3NC(=O)N(C3=O)C4CC[NH+](CC[NH+](C)C)CC4 (**I10**) | 462.4 | 2 | 7 | 71.68 | 6 |

\# MW : Molecular Weight

\# HBD : Hydrogen Bond Donors

#HBA : Hydrogen Bond Acceptors

\# TPSA : Topological Polar Surface Area

#I1….I10 : IDs given for top 10 derivatives.

**Table S9**: Depicts descriptor values of derivative compounds.

| Compound ID | D1 | D2 | D3 |
| --- | --- | --- | --- |
| I1 | 151.1166059 | 2.82462 | 6.293419279 |
| I2 | 151.3974003 | 2.68199 | 7.883446354 |
| I3 | 148.4421265 | 2.93142 | 7.252053952 |
| I4 | 150.0656818 | 2.76036 | 6.555356892 |
| I5 | 152.4493997 | 2.91072 | 7.100851909 |
| I6 | 151.0196726 | 2.78412 | 7.9490915 |
| I7 | 148.6830467 | 2.82462 | 7.239214974 |
| I8 | 150.6195159 | 2.82462 | 7.469083885 |
| I9 | 150.3058643 | 2.82462 | 7.469083885 |
| I10 | 151.0924563 | 2.92039 | 7.618251098 |
